# Flow alters fibrin molecular and network structure and decreases binding of fibrinolytic enzymes

**DOI:** 10.64898/2026.07.24.740662

**Authors:** Sajjad Norouzi, Ying Chou, Kashvika T. Ratheesh, Hieu Duc Tran, Yui Zhu, Xinhe Wang, Anh-Thu Nguyen, Farkhad Maksudov, Jan N. Fuhg, Manuel K. Rausch, Hsin-Chih Yeh, Kenneth A. Marx, Valeri Barsegov, Sapun H. Parekh

## Abstract

Thrombolysis with tissue Plasminogen Activator (tPA), approved for treating acute ischemic stroke (AIS) within 3–4.5 h of symptom onset, converts Plasminogen (Plg) to Plasmin to degrade fibrin, while fibrin also enhances Plg activation by binding to both tPA and Plg. Therefore, in this study, we investigated whether arterial-like flow, characteristic of AIS, alters fibrin structure and susceptibility to fibrinolysis. Using a newly developed platform for quantitative imaging and spectroscopy, we found that flow generates denser fibrin networks with reduced molecular transport despite reduced protofibril packing within fibrin fibers. Raman spectroscopy revealed an α-helix-to-β-sheet transition, accompanied by reduced Plg and tPA binding, although the reduction in tPA binding emerged only after prolonged flow exposure. Consistently, multi-scale molecular dynamics simulations showed that the Plg binding site destabilized at lower forces than the primary tPA binding site. Together, these multi-scale findings help explain the limited efficacy and narrow therapeutic window of thrombolysis.

## Introduction

Thrombolytic therapy is an approved treatment for acute ischemic stroke, with a therapeutic window of 3−4.5 hours after symptom onset^1,2^. It involves the conversion of Plasminogen (Plg) to Plasmin (Pln) by the serine protease enzyme, tissue Plasminogen Activator (tPA), followed by fibrin (Fn) degradation. However, this therapy remains limited in several respects. Even within the recommended treatment window, only ∼33% of patients achieve favorable outcomes following tPA administration^3^. Additionally, every 30-minute delay in treatment initiation reduces the probability of a favorable clinical outcome by ∼ 15%^4^. The limited efficacy of thrombolytic therapy and the associated risk of hemorrhagic complications therefore remain major clinical challenges^4–6^.

Improving the transport and delivery of tPA to the thrombus may appear to be the most obvious solution. However, several studies have already shown that tPA can effectively reach the thrombus during intravenous thrombolysis^7^. Moreover, retrieved thrombi exposed *ex vivo* to high concentrations of Plg and tPA (with no transport problem) still remain partially or completely resistant to lysis after 1 hour^8^. These observations indicate that delivery alone is not sufficient to explain thrombolytic resistance.

Another factor is thrombus composition. Thrombi contain red blood cells (RBCs), platelets, fibrin fibers, neutrophils, and other components^9^. Platelet- and fibrin-rich thrombi are more common in arteries and are generally more resistant to lysis than RBC-rich thrombi found in veins^10^. These compositional differences arise largely from the distinct hemodynamic environments in arteries and veins. Arteries experience higher shear rates and shear stresses than veins. Shear rates increase from ∼10 s^-1^ in veins to ∼2000 s^-1^ in small arteries (shear stress: 0.35−70 dynes/cm^2^)^11^, resulting in arterial and venous thrombi with distinct mechanisms of formation, compositions, and structural properties^9,12,13^. Consistent with these differences, fibrinolysis by tPA is slower *in vitro* for clots formed under arterial shear conditions than for those formed under venous flow^14^. These observations suggest that blood flow influences the properties of blood clots and thrombi and may directly affect the response to thrombolytic therapy.

TPA-mediated thrombolysis relies on fibrin to play two interconnected yet independent roles: (1) fibrin acts as a binding substrate for both Plg and tPA, brings them into spatial proximity, and consequently enhances Pln generation, and (2) fibrin serves as the substrate targeted and degraded by Pln. Plg binds to fibrin’s Aα148−160 through its Kringle domains and tPA binds primarily to fibrin’s γ312−324 via its finger domain. Alterations in fibrin molecular structure, particularly at and around these binding sites, are expected to influence the Plg and tPA binding and ultimately impair fibrinolysis efficiency. Such structural changes can arise from mechanically induced fibrin unfolding^15^, fibrinogen mutations^16^, or competition from other molecules for the Plg and tPA binding sites^17^, potentially affecting both fibrin’s functional roles. In the present study, we specifically focus on fibrin’s first role as a binding substrate for Plg and tPA. This role represents the first essential step in thrombolysis, as fibrin-bound Plg and tPA promote efficient Pln generation. We therefore investigated the role of hydrodynamic forces generated by flow in altering fibrin structure and, consequently, affecting fibrin ability to support Plg and tPA binding.

To study the response of fibrin to flow in isolation, other blood components must be excluded, and only fibrinogen (Fg; precursor of fibrin) should be used. Several experimental platforms have been developed to form blood or plasma clots under flow, including Badimon chambers, parallel-plate flow chambers, and microfluidic channels^18–22^. These systems typically rely on activation of the intrinsic or extrinsic coagulation pathways in whole blood or plasma through prothrombotic surfaces coated e.g., with collagen or tissue factor. However, these systems are not suitable for studying fibrin’s isolated response, as numerous proteins, lipids, and cells in blood or plasma can interact with fibrin and influence its behavior under flow. More importantly, none of these activating substrates can directly and independently convert fibrinogen to fibrin.

Physiologically, fibrinogen is activated by the serine protease, thrombin, through enzymatic cleavage of fibrinopeptides A and B^23^. The resulting fibrin monomers first polymerize longitudinally and then aggregate laterally to form fibrin fibers, which branch out, ultimately producing a homogeneous and isotropic fibrin network in three-dimensional space. However, to form fibrin under flow, direct addition of thrombin to flowing fibrinogen leads to rapid bulk fibrin formation throughout the solution, which can obstruct the flow path and fail to recapitulate the heterogeneous and anisotropic structure of fibrin in blood thrombi *in vivo*^9^. To overcome these limitations, Neeves et al. developed a microfluidic system comprising two perpendicular channels separated by a membrane^24^. In this design, fibrinogen is perfused through one channel while thrombin diffuses from the top channel through the membrane, initiating fibrin formation at the interface. Although effective, this platform is relatively complex and costly to fabricate and is not readily compatible with structural characterization methods such as Raman spectroscopy due to potential spectral interference from the device materials.

To address these limitations, we developed a simpler approach. Rather than delivering thrombin through a separate channel, we coated the coverslip with thrombin and flowed fibrinogen over it. Combined with a parallel-plate flow chamber, this approach enabled fibrin formation under flow while remaining compatible with structural analyses and time-dependent studies of fibrin remodeling. Furthermore, we coupled these *in vitro* measurements with *in silico* experiments on fibrin monomers and oligomers, in which we mimicked hydrodynamic flow conditions by subjecting fibrin molecules to pulling forces that span the experimentally relevant magnitude and direction ranges. This powerful combination of experiments and computer simulations has helped us understand experimental results on the biomolecular interactions between fibrin and the Plg and tPA enzymes.

## Materials and Methods

### Fibrinogen solution and buffer preparation

Fibrinogen was purchased from Enzyme Research Laboratories (FIB1), thawed at 37 °C, aliquoted, and stored at -80 °C until use. Aliquots were thawed each time at 37 °C and diluted in Tricine buffer (10 mM Tricine (T0377, Sigma Aldrich), 150 mM NaCl (S271-3, Fisher Scientific), 3 mM CaCl_2_ (746495, Sigma Aldrich), and 0.005% Tween 20 (P7949, Sigma Aldrich), pH 7.4) supplemented with 0.25% (w/v) Methylcellulose (MC; 036718.A4, ThermoFisher). MC was added to adjust the solution viscosity to approximate that of blood (∼3 cP^25,26^) (see supplementary Materials and methods and **Fig.S1**). For fluorescence imaging of fibrin network structure and morphology, fibrinogen solutions were supplemented with Alexa Fluor 647-labeled fibrinogen (F35200, ThermoFisher) at a 1:200 molar ratio of labeled to unlabeled protein in all experiments unless otherwise noted.

### Fibrin formation under flow

To form fibrin fibers under flow, thrombin was immobilized directly onto the glass surface. Human α-thrombin was purchased from Enzyme Research Laboratories (HT 1002a), aliquoted, and stored at -80 °C until use. Prior to fibrin formation, thrombin was diluted to 10 U/mL in Tricine + 0.25% MC buffer and applied to a defined 2 × 2 mm region on a #1 (24 × 60 mm) cleaned glass coverslip. Thrombin solutions were used either freshly prepared or stored at 4°C for up to 3−4 days prior to use. Coverslips were cleaned by sonication in 2% Micro-90 solution for 20 minutes, stored in a clean glass box, and plasma-treated for 1.5 minutes immediately before use. The coating region was confined using hydrophobic ink drawn with the narrow end of a 10 μL pipette tip. Thrombin solution (<2 μL) was incubated on the surface for 15 minutes at room temperature to promote adsorption; additional solution was added as needed to prevent drying during incubation. The surface was then washed 3−5 times with deionized water and then air dried (**Fig.1a-i)**. Upon incubation of fibrinogen solution on the thrombin-coated region, a uniform fibrin gel consistently forms within the solution. (**Fig.1a-ii,iii**).

This approach was implemented in a parallel-plate flow chamber. Parallel-plate flow chambers consist of rectangular cross-section channels that support well-defined laminar flow and are widely used for controlled shear experiments^20,27^. To fabricate the mold, a #1 (24 × 40 × ∼0.145 mm) glass coverslip was cut into a 24 × 3.6 mm strip and adhered to a 1 mm glass slide using a thin layer of glass glue. The assembled molds were placed in a custom 3D-printed enclosure, treated with a hydrophobic coating ((Tridecafluoro-1,1,2,2-tetrahydrooctyl) Trichlorosilane; SIT8174.0, Gelest), and reused across experiments. Polydimethylsiloxane (PDMS; Sylgard 184, 10:1 base-to-curing-agent ratio) was then poured over the mold and cured at 65 °C for 1 hour. After curing, the PDMS slab was peeled from the mold, forming the channel structure. Inlet and outlet ports were created at the distal ends of the channel using a 1.2 mm biopsy punch (504530, World Precision Instruments). For device assembly, the coverslip was first coated with thrombin (10 U/mL) for 15 minutes (**Fig.1b-i**) and then air-dried. The coated region was protected with a small coverslip fragment to prevent exposure and potential denaturation during plasma treatment. Both the PDMS channel and the coverslip were then plasma-treated for 1 minute and brought into conformal contact to form an irreversible bond. Immediately prior to bonding, the protective small coverslip was removed, and the PDMS channel was aligned such that the thrombin-coated region was positioned at the center of the flow path (**Fig.1b-ii**). The inlet and outlet ports were then connected to a peristaltic pump (15124696, Masterflex™ or LS0216-2, MSE Supplies). The channel was initially perfused with Tricine + 0.25% MC buffer to hydrate the channel and remove trapped air (**Fig.1b-iii).** The inlet tube was then inserted into a tube containing fibrinogen solution maintained at 37 °C in a water bath. After 2−3 minutes of initial perfusion, the flow was paused for 10 minutes. This pause allowed initial fibrin molecules to anchor to the coverslip and promote subsequent fibrin formation under flow over longer time. Without it, no fibrin fibers were observed under flow for up to 1 hour. Subsequently, the flow rate was set to 1.1 mL/min, corresponding to a shear rate of 1500 s^-1^ and a wall shear stress of 4.6 Pa (for a laminar Newtonian fluid, see supplementary Materials and Methods)^11^, within the physiological range of arterial shear^11^. Because both blood and MC-based solutions exhibit shear-thinning behavior^26,28^, the calculated shear stress is just an estimate, assuming a Newtonian fluid. Perfusion was carried out for 1 or 3 hours under closed-loop conditions, allowing fibrinogen to interact with the thrombin-coated surface and form fibrin fibers within the chamber (**Fig.1b-iii**). All other perfusion steps, including loading, washing, buffer exchange, and post-gel enzyme exposure, were performed at 0.5 mL/min in all experiments. For static conditions (no flow), the chamber was first filled with fibrinogen solution. The tubing was then sealed, and the entire device was incubated at 37 °C for 1 hour.

Remaining (unpolymerized) fibrinogen concentration in the reservoir was quantified by absorbance at 280 nm using an Eppendorf BioSpectrometer and the fibrinogen extinction coefficient (extinction coefficient (1%) = 15.1). When fluorescent fibrinogen variants were used, interference from dyes was corrected using their absorbance at the maximum excitation wavelength. For static clots, 10 µL of solution was collected from the chamber after removing one of the tubes and measured for absorbance at 280 nm to calculate the unpolymerized fibrinogen concentration.

### Fluorescent recovery after photobleaching (FRAP)

FRAP experiment was performed using Oregon Green dextran (70 kDa; D7172, ThermoFisher) to approximate the molecular weight of tPA. After fibrin formation ([Fg]=1 mg/mL), the channel was washed with buffer and perfused with 1 μM dextran in phosphate-buffered saline (PBS; 70011044, ThermoFisher). The sample was incubated for 1 hour at room temperature to allow dextran to diffuse throughout the fibrin network. The channel was then sealed and imaged on the confocal microscope with a 30×, 1.05 NA silicone oil immersion objective (UPLSAPO30XS, Olympus Corporation). For each experiment, dextran recovery was measured both in the fibrin gel and in the surrounding solution with minimal fibrin. A circular region of interest (ROI) with an 11 µm radius was used for photobleaching in each frame across all the experiments. For in-gel measurements, fibrin fibers were identified using either 640 nm excitation at 0.2% laser power (detector HV: 500) or 560 nm excitation at 1.3% laser power (detector HV: 500). Photobleaching was performed with a 488 nm laser at 95% power for 10 seconds. Imaging then resumed at 3% laser power (detector HV: 650) with a time interval of 0.88 s. Images were acquired at an optical zoom of 3 with a 256 × 256 pixel per frame, two frame-averaging, and a dwell time of 2 µs. Fluorescence recovery was recorded for approximately 2.5 minutes. The emission windows for the 488 nm and 640 nm lasers were set to 500−540 nm and 650−750 nm, respectively.

The fluorescent intensity in each ROI was corrected for photobleaching and normalized to the range [0, 1]^29^. Photobleaching correction was performed using pixels located more than 50 µm from the ROI center, as they were minimally affected by ROI photobleaching. The corrected recovery intensity was plotted as a function of time and fitted with a double-exponential function to determine the half-recovery time (t_1/2_), defined as the time at which the fitted curve reaches half of the final plateau value. The diffusion coefficient was calculated using 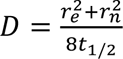 , where r_n_ is the ROI radius (11 µm) and r_e_ is the effective radius^30^. To find r_e_, the first post-bleach frame was normalized to the pre-bleach image to obtain the photobleaching profile. The image was then radially binned (0.5 µm bin size), and the mean intensity was calculated within concentric rings up to 50 µm from the ROI center. The resulting radial intensity profile was fitted with a Gaussian function. The effective radius, r_e_, was defined as the distance at which the fitted Gaussian decayed to 14% of the difference between the peak (center) and baseline (far-field) intensities.

### Fluorescence lifetime imaging microscopy (FLIM) and analysis

To compare intermolecular packing within fibrin fibers among different conditions, we used FAM (5,6)-isothiocyanate (FITC; 3524, Lumiprobe) as a self-quenching fluorescent tracer^31^ to covalently label fibrinogen. FAM (5,6)-FITC was dissolved in dimethyl sulfoxide (DMSO) to a stock concentration of 2 mM. A 5 mg/mL fibrinogen solution was then mixed with FAM (5,6)- FITC at a molar ratio of 1:4 (Fg:FITC) and incubated for 1 hour at room temperature on an orbital shaker. Excess unbound dyes were removed using Pierce™ Dye Removal Columns (22858, ThermoFisher). The final fibrinogen concentration and degree of labeling in the stock solution were 4.6 mg/mL and 0.97 FITC molecules per fibrinogen molecule, respectively. These values were determined from absorbance measurements at 280 nm and 492 nm using an Eppendorf BioSpectrometer and the known extinction coefficients of fibrinogen and FITC. All static and flow experiments were performed using 1 mg/mL fibrinogen (of which 5% was FITC-labeled) and details in the previous sections.

FLIM measurements were performed using an ISS Alba 5 laser scanning confocal system built around a Nikon Eclipse TE2000-U microscope and a 60× water-immersion objective (Nikon Plan Apo, NA = 1.2). The emitted fluorescence was collected by an avalanche photodiode detector (SPCM-AQR-15, PerkinElmer) and processed using the FastFLIM frequency-domain acquisition board (ISS Inc.). Fluorescence lifetime data were acquired and analyzed using the VistaVision software (ISS Inc.). Before data acquisition, the instrument response function (IRF) was calibrated using 1−6 µM fluorescein (L13251.36, ThermoFisher Scientific) in PBS with a known lifetime of 4.0 ns^32–34^. The sample was excited at 488 nm using a laser diode (LDH-D-C-488, PicoQuant) at 40 MHz modulating frequency, and the emitted fluorescence was collected through a 200 µm pinhole and filtered using a band-pass emission filter (FF02-525/40-25, Semrock). Each frame consisted of 256 × 256 pixels with a 50 µs pixel dwell time. To acquire enough photon counts for lifetime analysis, pixels from 6−8 consecutive frames were pooled into 1 image for 1 field-of-view (FOV).

For frequency-domain FLIM analysis, the phasor plot analysis was used^35,36^, where each pixel was mapped to its corresponding phasor coordinates (g, s) and displayed on the universal semicircle of the phasor plot. All pixels from the pooled image will create a phasor distribution.

For single-lifetime species or single exponential decay species, the phase lifetime, r_p_ , and modulation lifetime, r_m_, are calculated using the following equations:

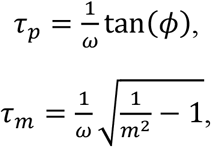

where ф is the phase, the angle between the center of the phasor distribution and the x-axis, *m* is the module, the distance from the center of the phasor distribution and (0,0), and ω is the modulation frequency of the excitation laser, which is 40 MHz.

In this study, a Gaussian and a median smoothing filter with a bin width of 1 pixel were applied to the phasor distribution. Approximately 10% of the darkest pixels were excluded from the lifetime analysis. All the fluorescence lifetimes reported in this study correspond to phase lifetimes (r_p_). Within a sample, different FOVs were taken. The mean r_p_ of 1 FOV was considered as a single technical replicate, and multiple samples of the same condition were treated as biological replicate.

### Broadband coherent anti-Stokes Raman scattering (BCARS) microscopy and analysis

To study the fibrin molecular structure, we used the BCARS hyperspectral microscopy, previously used for fibrin, collagen, and protein condensate studies in our lab^37–41^. For BCARS measurements, the PDMS chip was fabricated with reduced thickness above the channel center. This allowed the collection objective to approach the excitation objective without contacting the chip for proper focusing. After formation, the fibrin gel was washed with Tricine buffer (without MC). This step removed residual MC from the surrounding solution and minimized its interference with the fibrin Raman signal.

A 1064 nm laser with sub-100 picosecond pulses and 1 MHz frequency, and a broadband 1100- 2400 nm supercontinuum laser (Leukos Opera HP, Leukos) were used as pump and Stokes beams, respectively. The two beams were focused through a 100×, 0.85 NA objective (LCPLN100XIR, Olympus Corporation), and the transmitted signal was collected using a 20×, 0.4 NA objective (M- 20X, Newport Corporation). Pixel sizes of 0.5 µm or 1 µm were used, and the CCD integration time was adjusted to maximize signal while avoiding saturation.

BCARS raw signals were converted into Raman spectra following established methods^42,43^. Spectra were acquired over a Raman shift range of 700−4000 cm^-1^ with an average spectral resolution of ∼4 cm^-1^. After phase retrieval, spectra were normalized to the total Amide I band area (1630−1700 cm^-1^) and background-subtracted using the intensity at the lowest Raman shift (∼700 cm^-1^). This normalization assumes that the overall protein secondary structure, as reflected by the Amide I band, is preserved across conditions. Each spectrum represents the average of a 3 × 3 pixel region. To compare full Raman spectra across different groups, dimensionality was reduced to two components using principal component analysis (PCA) in MATLAB. Only spectral regions with Raman shifts below 1700 cm^-1^ and above 2800 cm^-1^ were included, as biological samples show minimal Raman signal in the intermediate range^44^.

For secondary structure analysis of fibrin, the Amide I region was fitted with seven Lorentzian curves^45^. Each curve represents a specific secondary structural motif, and the sum of their areas accounted for the total Raman intensity in the Amide I region. The full width at half maximum (FWHM) values were initially optimized to obtain α-helix and β-sheet contents of ∼31% and ∼37%, respectively, within the range of previously reported values^46–48^. The optimized FWHM values were then fixed, and together with constrained peak-position ranges, were used in subsequent curve fittings (**Table S1**). The percentage of each secondary structural motif was calculated as the area under the corresponding fitted component divided by the total area of all fitted secondary structure components.

### Structural models of fibrin polymers

#### Atomic structure of human fibrin monomer

The atomic structural model of human fibrin was derived from the structure of fibrinogen monomer available from the Protein Data Bank (PDB ID entry 3GHG)^49,50^. This model contains resolved residues Aα27–200, Bβ45–461, and γ1–394, and lacks the residues of αC region (residues Aα221–610). We used our previously developed fibrin monomer model comprising (residues Aα1–210, Bβ1–451, and γ1–411) in which the missing N-terminal segments (residues Aα1–26 and Bβ1–44) and the disordered C-terminal γ-tail (residues γ395–411) were computationally reconstructed as described elsewhere^51^. To reconstruct the missing C-terminal segment of the Aα chain (residues Aα211–610), which includes the αC region (Aα221−610) and is absent from all experimentally resolved fibrin and fibrinogen structures, the sequence spanning residues Aα211−610 was used as the input for AlphaFold 3.0^52^.

The reconstructed fibrin fragment was subsequently appended to residue Aα210 using the VMD 1.9 software^51^. This yielded a full-length atomic structure of human fibrinogen comprising 2,904 amino acid residues, including two Aα1–610, two Bβ1–461, and two γ1–411 fragments. To arrive at the atomic model of human fibrin, the N-terminal fibrinopeptides A (residues Aα1–16) and fibrinopeptides B (residues Bβ1–14) were removed (**Fig.3d**). The following fibrin monomer systems were used in the simulations: (1) the half-fibrin monomer with one of the two αC regions (half-Fn; **Fig.3e**), which contains residues Aα17–610, one of the two β-nodules (residues Bβ15– 461), and one of the two γ-nodules (residues γ1–411); and (2) the half-fibrin monomer without the αC region (half-Fn/des-αC; **Fig.5a**), containing residues Aα17–210, residues Bβ15–461, and residues γ1–411, but excluding the residues connecting to and αC (Aα211-610). *Coarse-grained model of fibrin oligomers (FO)*: We also used (3) the Cα-atom based structure of short double-stranded fibrin oligomer (FO4/4) containing 4/4 fibrin monomers in the first/second strands (**Fig.S2c**), reconstructed as described in our previous study^51^. This full-length fibrin structure has the αC chains, the A:a and B:b knob-hole bonds, and the γ-γ crosslinking sites. This structure has the αC chains, the A:a and B:b knob-hole bonds, and the γ-γ crosslinking sites.

**Figure 1.**
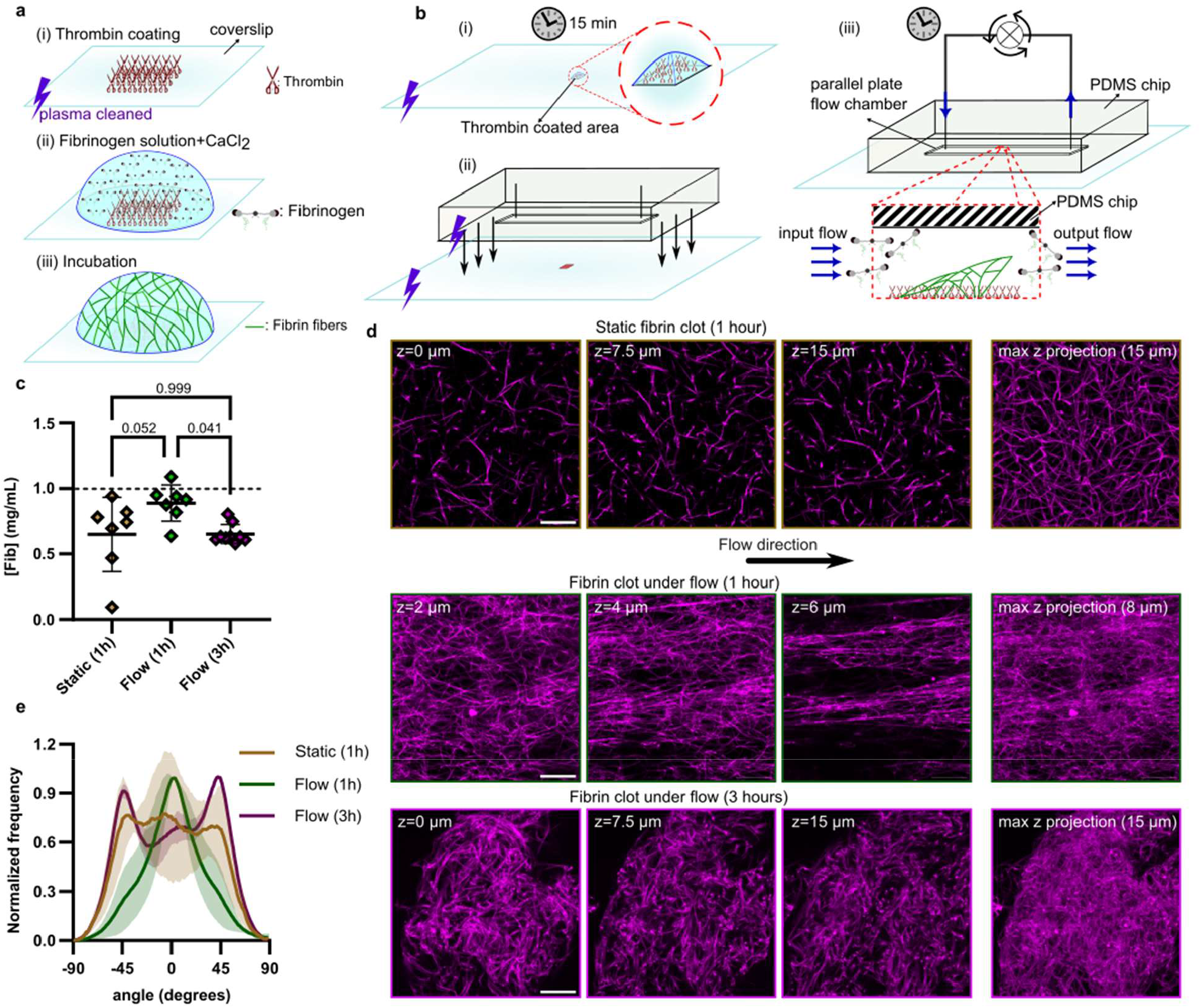
Characterization of fibrin fibers formed under flow. **a)** Schematic of fibrin fibers form on a thrombin-coated surface. Thrombin is immobilized on a plasma-cleaned coverslip (i), which is then surrounded by fibrinogen solution in the presence of CaCl₂ (ii), leading to fibrin formation over time upon incubation (iii). **b)** Fibrin formation on the thrombin-coated surface is integrated with a parallel-plate flow chamber. After thrombin incubation (i), the coverslip and PDMS chip are plasma-cleaned and bonded such that the coated region is positioned within the channel (ii). The inlet and outlet are then connected to a peristaltic pump to establish flow and enable fibrin formation (iii). **c)** Fibrinogen concentration remaining in the reservoir after flow experiments. **d)** Fluorescence images of fibrin fibers at different z-planes and their corresponding maximum-intensity projections (scale bar= 20 μm). The intensity contrast was corrected individually for better visualization. **e)** Normalized orientation distribution of fibrin fibers under different conditions, showing a transition from isotropic organization under static conditions to increased alignment under flow. The lines show the mean value and standard deviation is shown as the shaded area (n=3).

**Figure 2.**
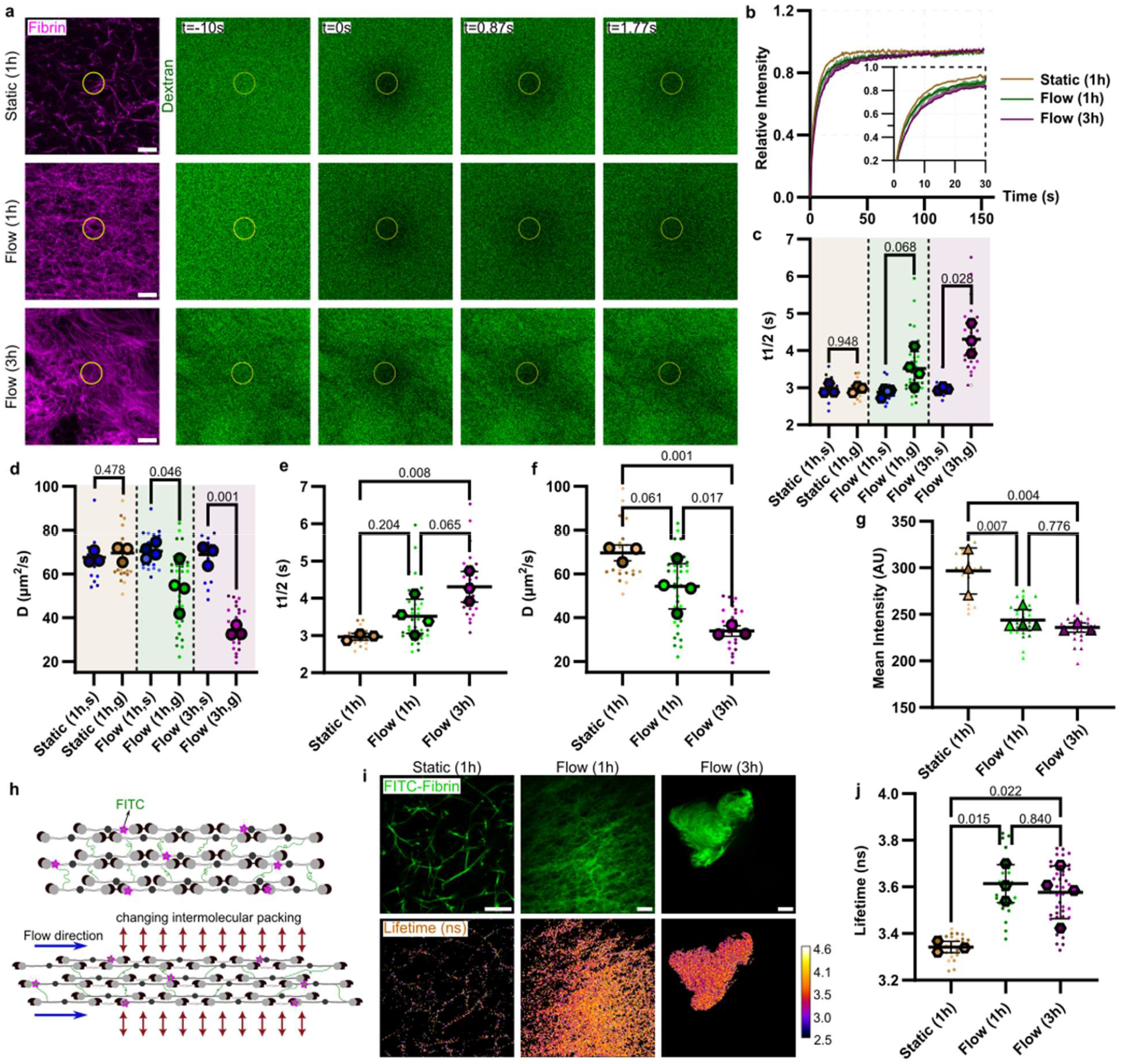
Flow produces a denser fibrin network with reduced molecular accessibility and altered intermolecular packing. **a)** Fluorescence images of fibrin clots before and after photobleaching under different conditions. Scale bar = 20 μm. Contrast was adjusted independently for each group. **b)** Photobleaching-corrected fluorescence intensity within the ROI as a function of time; inset shows a magnified view of the recovery region. The shaded area represents ±1 standard deviation around each curve (n = 3, 4, and 3 for Static (1h), Flow (1h), and Flow (3h), respectively). **c)** Comparison of the recovery half-time, (t_1/2_), for dextran measured in the gel (g) and solution (s) within the same flow chamber. **d)** Comparison of dextran diffusion coefficients in the gel and solution across conditions. **e)** Comparison of in-gel (t_1/2_) values across conditions, showing slower diffusion in fibrin formed under flow, particularly after 3 hours. **f)** Comparison of in-gel diffusion coefficients across conditions. **g)** Dextran intensity within fibrin-masked regions, demonstrating reduced accessibility of dextran to fibrin fibers formed under flow. **h)** Schematic of FITC-labeled fibrin molecules within a fiber, illustrating how changes in intermolecular packing may alter fluorophore proximity and fluorescence lifetime. **i)** Representative images of FITC-labeled fibrin fibers and their corresponding fluorescence lifetime maps. Scale bar = 20 μm. **j)** Comparison of FITC lifetimes across conditions, showing increased lifetime in fibrin formed under flow. In all plots, each large circle represents one biological replicate (the mean of technical replicates), while smaller circles of the same color represent individual technical replicates.

**Figure 3.**
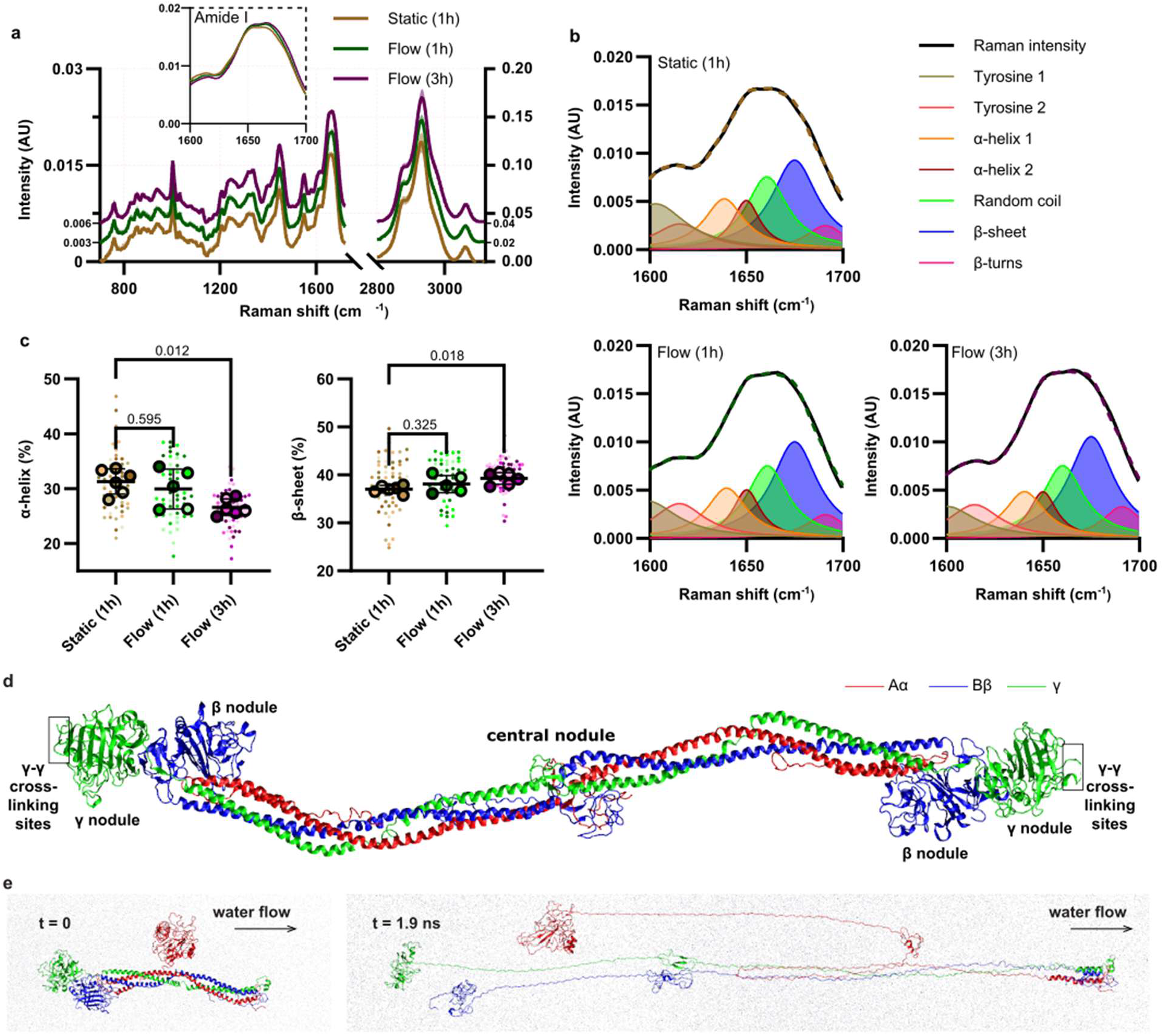
Flow alters the structure of fibrin. **a)** Raman spectrum of fibrin under different conditions, shown for Raman shifts <1700 cm⁻¹ (left axis) and >2800 cm⁻¹ (right axis). The intermediate region is excluded due to the absence of characteristic peaks for biological materials. Spectra for flow (1h) and flow (3h) are vertically offset for clarity; the magnitude of the offset is indicated next to each curve. The inset shows a zoomed-in view of the Amide I region. Bold lines represent mean spectra, and shaded regions indicate standard deviation. **b**) Lorentzian fits of the Amide I region used to extract secondary structural components. The curves correspond to fits of the mean Raman spectra, and the dashed line represents the sum of all fitted component curves. **c**) Quantitative comparison of α-helix and β-sheet content derived from **b**, showing a transition from α-helix to β-sheet structure. Each large circle represents one biological replicate, while smaller circles (matching color) denote individual technical replicates. **d**) Structure of human fibrin monomer showing the central nodule, the γ-nodules, the β-nodules, and the γ-γ-crosslinking sites (the αC domain and αC connector are not shown). **e**) Snapshots of half-fibrin fragment (half-Fn) before (left panel) and after (right panel) the water flow is switched on, showing the flow induced unfolding of half-Fn.

**Figure 4.**
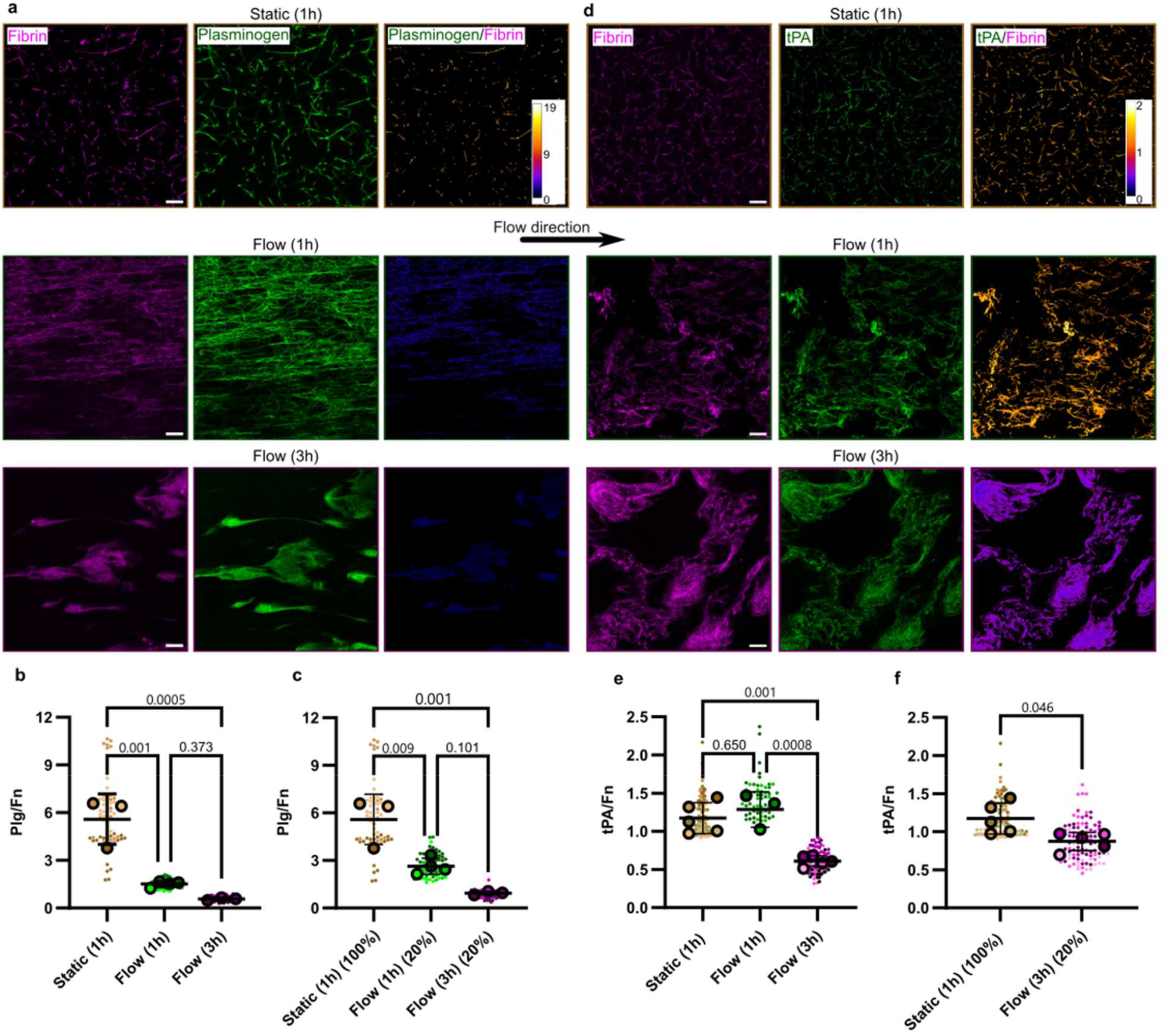
Flow reduces the Plasminogen and tPA binding to fibrin fibers. **a**) Representative fluorescence images of fibrin fibers formed under different conditions and subsequently incubated with Plg, along with corresponding Plg/Fn intensity ratio maps. **b**) Quantitative analysis of the Plg/Fn intensity ratio across conditions, showing reduced Plg binding in fibrin formed under flow. Small circles represent the mean Plg/Fn ratio for each frame, while larger circles (matching color) indicate the mean value across frames for each experiment. **c**) Comparison between flow conditions, where only the top 20% of pixels with the highest Plg/Fn ratios are included, and static condition, where 100% of pixels are analyzed. This comparison aims to reduce the transport and availability limitations that could otherwise artifactually reduce the Plg/Fn ratio in flow conditions. **d**) Representative fluorescence images of fibrin fibers incubated with labeled tPA after formation, along with corresponding tPA/Fib intensity ratio maps (range: 0-2). **e** Quantitative analysis of the tPA/Fn intensity ratio across conditions. Small circles represent the mean value in each frame while their combined average in each experiment is shown by large circles (matching color). **f**) Comparison between the Static (1h) and Flow (3h) tPA/Fn ratio where only 20% of pixels with the largest ratios are included for Flow (3h). All fibrin, Plg, and tPA channels were independently contrast-adjusted for visualization, while ratio maps were displayed using a consistent contrast scale in each Plg and tPA groups. The scale bar is 20 μm.

**Figure 5.**
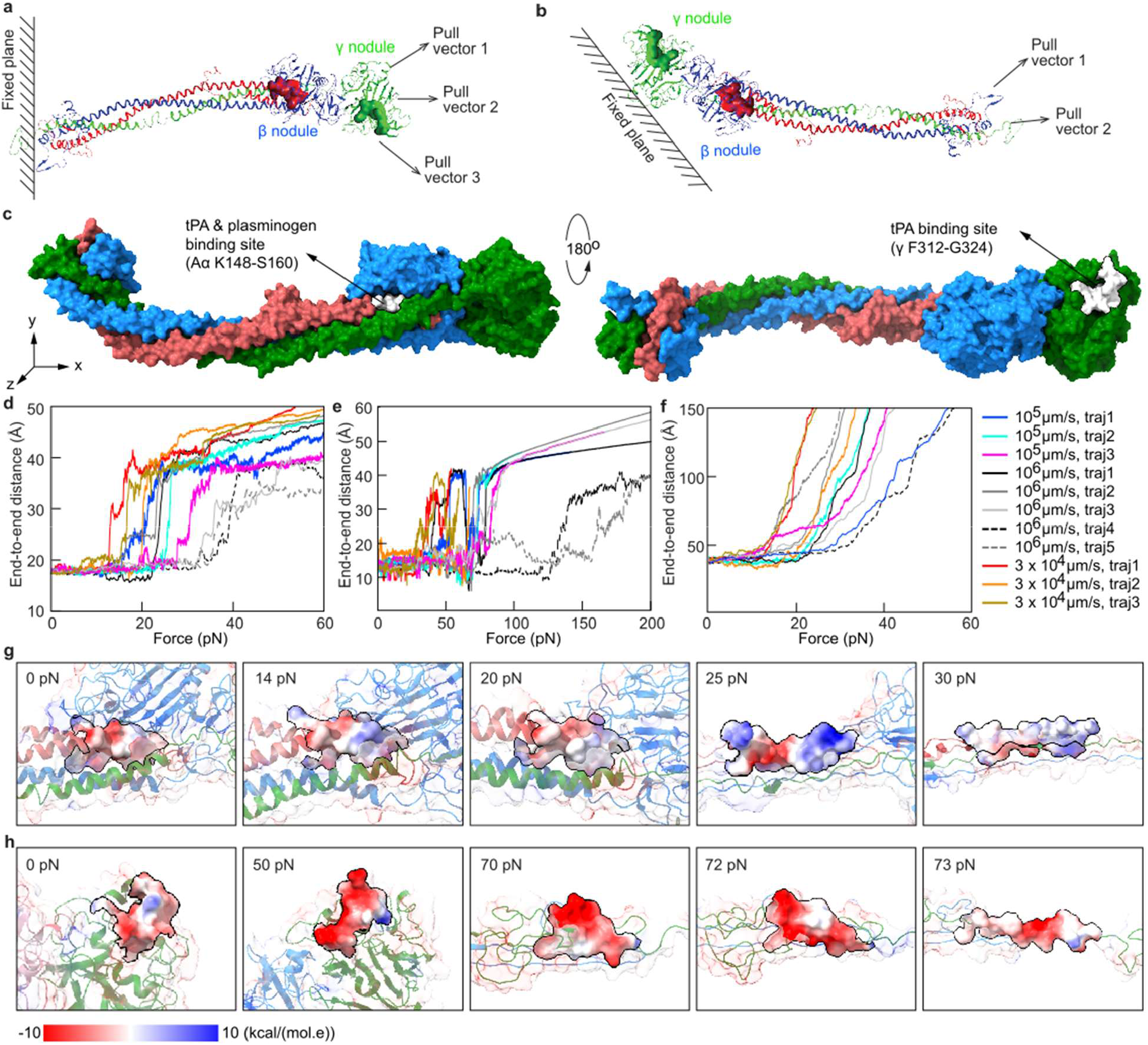
Structural changes in fibrin tPA- and Plasminogen-binding sites during force unfolding in silico. **a**) Schematic of force conditions illustrated for Half-Fn/des-αC model implemented in the ramped force simulations to trigger the unfolding transitions. The pulling forces mimicking hydrodynamic flow are applied to the β- and γ-nodules along three directions relative to the fibrin longitudinal axis, while the central E region is constrained. **b**) Alternative pulling configuration in which forces were applied to the central E region while the β- and γ-nodules are constrained. c) Molecular surface representation of half-Fn/des-αC with the Aα, Bβ, and γ chains colored in red, blue, and green, respectively. In panels a-c, the color denotation is same as in Figs.3d,e. The tPA/Plasminogen-binding region in the Aα chain (residues K148–S160) and the tPA-binding site in the γ chain (residues F312–G324) are highlighted in white. d) Distance between the C_α_-atoms spanning the Aα residues K148–S160 (size of the binding site for tPA and Plasminogen) as a function of applied force. **e**) Distance between the C_α_-atoms spanning the γ chain residues F312– G324 (size of the binding site tPA) as a function of force. **f**) Distance between the center of mass of K148-S160 and F312-G324 as a function of applied force. **g**) Electrostatic surface representation of the binding site for tPA and Plasminogen on Aα-chain binding site (residues K148-S160) at various force levels (in the 0–30-pN range). **h**) Electrostatic surface representation of the binding site for tPA on γ-chain (residues F312–G324) at various force levels (in the 0–73 pN range). Surface colors represent electrostatic potential, with red indicating negative potential and blue indicating positive potential.

### Molecular Dynamics simulations

#### Implicit solvent modeling

We modeled the full-length fibrin molecule with the αC chain (full-Fn) and the half fibrin molecule without the αC chain (half-Fn/des-αC) using the Solvent Accessible Surface Area (SASA) model of implicit solvation in conjunction with CHARMM19^51,53,54^. *Explicit solvent modeling:* We modeled the half-fibrin monomer with the αC chains (half-Fn) using an explicit solvent model in conjunction with the ff19SB force field^55^ implemented in the Amber 2020 software^56^ patched with the PLUMED plugin^57^. The atomic structure of half-Fn system was solvated in a rectangular water box with ∼564,971 water molecules (∼1897 nm^3^ solvation box). Counterions were added to neutralize the system, and Na⁺ and Cl⁻ ions were included to achieve a physiological concentration of 150 mM. Next, we performed energy minimization, heating, and equilibration. Due to differences in solvent representation, implicit- and explicit-solvent models followed distinct preparation protocols. For implicit-solvent systems (half-Fn/des-αC), energy minimization was performed for 1 ns. Systems were then heated to 300 K over 0.3 ns, followed by equilibration at 300 K for 10 ns. For the explicitly solvated half-Fn system, a multi-stage restrained minimization and equilibration protocol was applied. Energy minimization was performed in four stages: (1) 25,000 steps with positional restraints of 25 kcal/mol/Å2 on solute heavy atoms; (2) 25,000 steps with restraints reduced to 10 kcal/mol/Å^2^; (3) 500,000 steps with 1 kcal/mol/Å2 restraints; and (4) 400,000 steps without restraints. The system was heated to 300 K over 0.3 ns. Equilibration was performed in multiple stages with gradually reduced restraints: 0.2 ns (50 kcal/mol/Å^2^), 0.2 ns (25 kcal/mol/Å²), 0.4 ns (10 kcal/mol/Å^2^), 0.4 ns (1 kcal/mol/Å^2^), 0.6 ns (0.5 kcal/mol/Å^2^), followed by 1.6 ns without restraints. *Coarse-grained modeling:* The native topology-based Self-Organized Polymer (SOP) model, implemented in the SOP-GPU package^51,58–60^, was used to describe fibrin oligomer FO4/4. The SOP model has been successfully used to explore various large-scale biomolecular assemblies, including fibrin oligomer^51,61^, microtubule filaments^62,63^, virus shells, and cell organelles^64–66^. For the fibrin oligomer FO4/4 system, we performed minimization, heating, and equilibration steps as described in our prior study^67^. *Analysis of MD trajectories:* Molecular Dynamics trajectories were visualized using VMD 1.9 and ChimeraX 1.12 and analyzed using custom Python scripts. Secondary structure was assigned using the DSSP algorithm^68^.

### Dynamic force protocol

#### Explicit solvent simulations

To simulate the non-equilibrium dynamic flow conditions, we employed the MD simulations in explicit solvent. A constant external bias has been applied to the center-of-mass (COM) of the oxygen atoms in water molecules to generate a unidirectional water flow along the x-direction (**Fig.3d**). To maintain the structural orientation of the fibrin molecule in the fluid flow, harmonic restraints were applied to the Cα-atoms of the residues in the β-nodule and γ-nodule or to the central E domain of the fibrin molecule. The force protocol options, which specify the constrained residues and flow velocity, are summarized in the SI (**Table S2**). These restrained regions served as stationary reference points for quantifying solvent-induced forces. To accommodate large molecular elongation due to fibrin unraveling, an adaptive solvation box extension approach was employed. Briefly, when the unraveling portion of the fibrin protein approached the boundary of the simulation solution box, the simulation was paused, and a pre-equilibrated solvent box was appended to accommodate the resulting molecular elongation. The expanded system was then energy-minimized, heated, and equilibrated under strong positional restraints to preserve the protein’s non-equilibrium conformation. Next, restraints were reduced as appropriate, and the external bias on the solvent COM was re-applied to resume the flow-driven molecular extension (**Fig.3f**). *Implicit solvent simulations:* To mimic conditions of the fluid flow in the implicit solvent simulations, we constrained one part of the molecule and exerted a time-dependent pulling force, *f*(*t*) = *r*_ƒ_*t*, to another part of the molecule. We used the force-loading rate, *r*_ƒ_ = *k*_sp_*v*_ƒ_ , where *v*_ƒ_ is the pulling speed of a virtual cantilever and *k*_sp_ is the cantilever spring constant^53,54^. We set *k*_sp_ = 300 pN/nm and varied *v*_ƒ_ in the 1−10^6^-μm/s range (see **Table S3**). For half-Fn/des-αC system, two types of pulling force protocols were employed. In the first case study, the force was applied to the Cα-atoms of amino acids in the central E domain, while the Cα-atoms of amino acid residues in the β- and γ-nodules were constrained (**Fig.S2a**). In the second case study, selected Cα-atoms of residues in the central E domain were constrained, while the pulling force was applied to the Cα-atoms of amino acids in the β- and γ-nodules (**Fig.S2b**). *Coarse-grained simulations:* For the fibrin oligomers FO4/4, the pulling protocol was described before^67^. Briefly, one (left) end is constrained, and the force is applied to the other (right) end (**Fig.S2c**, **Table S3**). *Geometry of force application:* Multiple pulling geometries together with the 1−10^6^ μm/s pulling speed range were considered to sample distinct protein unfolding pathways corresponding to the conditions of flow-induced shear stress. The constrained residues were selected to represent the major force-transmission regions in fibrin, including longitudinal connections through the β- and γ-nodules. For example, for the half-Fn/des-αC system, asymmetric force conditions were imposed by constraining amino acid residues in one terminal region (subset of the Cα-atoms in Bβ and γ chains) and applying forces to the Cα-atoms of amino acids in the opposite terminus. Distinct residue selections spanning the Cα-atoms in the β- and γ-nodules and Cα region are detailed in the SI (**Table S3**).

### Plasminogen and tPA binding assays and imaging

FITC-labeled human Plasminogen was purchased from Abcam (AB92770), aliquoted, and stored at -80 °C until use. For the binding assessment, it was diluted to 125 nM in PBS. After fibrin formation under static or flow conditions ([Fg]= 3 mg/mL), the samples were washed with Tricine buffer + 0.25% MC. The Plasminogen solution was then perfused into the channel and incubated for 30 minutes at room temperature. The channel was subsequently washed again with buffer prior to imaging. Confocal fluorescence imaging was performed with a 30×, 1.05 NA silicone oil immersion objective. Imaging was conducted sequentially in the green (Plg) and far-red (Fn) channels to minimize crosstalk. Fibrin fibers were first imaged using 640 nm excitation at 0.5% laser power (detector HV: 490), with an emission window of 650−750 nm. Plg was then imaged using 488 nm excitation at 1.5% laser power (detector HV: 550), with an emission window of 500−600 nm. Each image consisted of 1024 × 1024 pixels, acquired at an optical zoom of 2, with a dwell time of 2 µs and 2 line-averaging.

Images were segmented based on the fibrin channel intensity using Otsu’s method in MATLAB. For each frame (technical replicate), the mean ratio of Plg intensity to fibrin intensity (Plg/Fn) was calculated across all segmented pixels. Biological replicates were the means of these ratio values across all frames for each experiment.

For tPA binding, single-chain tPA (Innovative Research, IHUTPA85SC100UG) and FITC-labeled tPA (Innovative Research, IHUTPASCFITC100UG) were aliquoted as received and stored at -80 °C until use. Because fibrin contains more tPA binding sites than Plg binding sites, we used a lower fibrinogen concentration ([Fg] = 1 mg/mL) and a higher tPA concentration (8.82 μM, 1:9 ratio of labeled to unlabeled) to ensure an excess of enzyme. Following overnight incubation at room temperature on a platform rocker, the channel was perfused and washed with buffer to remove unbound enzymes and then subjected to fluorescence imaging. All the optical settings are the same as for Plg imaging, except that the excitation power of the 488 nm laser was 1%. As in the Plg image analysis, images were segmented based on fibrin intensity, and the ratio of tPA intensity to fibrin intensity (tPA/Fn) was calculated.

### Statistical analysis

All statistical analyses were performed in GraphPad Prism using either Student’s t-test or ANOVA. Tukey’s test was used for multiple comparisons. The exact p-values between the compared groups are shown on each graph, with a significance threshold of p < 0.05.

## Results

### Flow alters fibrin fiber network structure

To initiate fibrin formation at a specific location in a microfluidic channel, we coated thrombin on an area to locally cleave fibrinogen and initiate fibrin fiber formation. Although thrombin is restricted to a small area, fibrin fibers form throughout the channel (**Fig.S3**). This behavior is observed under both static and flow conditions (**Fig.1a-ii, iii)**. In the absence of immobilized thrombin on the surface, no fibrin fibers formed during 3 hours of fibrinogen perfusion (**Fig.S4**). Since we are forming fibrin under shear, we used a 0.25% (wt/wt) of methyl cellulose (MC) as an additive to match the viscosity of blood in our polymerization buffer (see methods). To confirm wthat neither 0.25% MC nor immobilized thrombin affects the fibrin molecular structure, Raman spectra were compared for fibrin formed under two conditions: (1) in Tricine buffer on a coverslip with soluble thrombin and (2) in Tricine buffer + 0.25% MC in the flow chamber with surface-coated thrombin, bot h after 1 hour of incubation with no flow. The secondary structure content was quantified by Lorentzian fitting and compared between the two groups (**Fig.S5**). The spectral overlap and the absence of significant differences in secondary structure demonstrate that neither 0.25% MC nor the mode of fibrin formation altered the molecular structure of fibrin. These conditions did, however, affect the kinetics of fibrin formation. As expected, increasing viscosity with 0.25% MC slowed fibrin formation compared with Tricine buffer alone in turbidity measurements (**Fig.S6**, supplementary Materials and Methods). In addition, fibrin formation proceeded more rapidly when thrombin was present in solution than when it was confined to the surface.

Localized activation by thrombin resulted in less overall fibrin formation relative to the initial fibrinogen concentration (**Fig.1c**). Under recirculating, peristaltic flow, fibrin formation was initially slower, with only ∼10% of fibrinogen consumed after 1 hour compared with ∼35% fibrin consumption under static conditions. However, fibrin formation increased with continued flow, rising from ∼10% fibrinogen consumption after 1 hour to ∼35% after 3 hours. Despite lower early fibrin yield, flow conditions generated a substantially denser fibrin network, whereas static conditions led to a more porous network structure (**Fig.1d**). Under 1 hour of flow, fibrin fibers aligned predominantly parallel to the flow direction (see the Supplementary Materials and Methods for details of the orientation quantification). This alignment was expected to increase further after 3 hours; however, the dynamic coupling between network growth and flow progressively perturbed the local flow field and led to a transition of single dominant alignment peak at 1 hour to a split distribution at 3 hours, with two peaks at approximately ±45° (**Fig.1e**).

### Flow slows diffusion within the fibrin network and reduces fibrin accessibility

One key property of blood clots is their accessibility to circulating enzymes and therapeutic agents. This is particularly important in thrombolytic therapy, where tPA must penetrate the clot to reach its interior. To assess fibrin accessibility, we performed FRAP analysis using dextran molecules with a molecular weight similar to tPA (70 kDa). After allowing dextran to equilibrate in the flow chamber, an ROI within the fibrin gel was photobleached, and fluorescence recovery was monitored (**Fig.2a**). The photobleaching-corrected and normalized intensity profile was fit to a double-exponential curve to determine t_1/2_, defined as the time at which the fitted curve reaches half of its final plateau value (**Fig.2b**). FRAP measurements were also performed in regions of the chamber with minimal fibrin to represent “in-solution” conditions.

Under static conditions, fibrin gels showed t_1/2_ values similar to those in solution (**Fig.2c**), suggesting that the 70 kDa dextran diffused freely through the fibrin network. In contrast, fibrin formed under flow, particularly after 3 hours, exhibited larger t_1/2_ values, indicating slower diffusion of the dextran probe within the gel. The diffusion coefficient was calculated by fitting a Gaussian function to the radial intensity profile of the normalized first post-bleach frame to determine the effective radius of photobleached area (**Fig.S7**)^30^. The diffusion coefficient of dextran in solution was ∼ 69 µm^2^/s, within the range of reported values for Bovine Serum Albumin of similar molecular weight (∼62 µm^2^/s with ∼66 kDa molecular weight)^69,70^.

Consistent with t_1/2_ measurements, static fibrin gels after one hour (static (1h,g)) showed a similar dextran diffusion coefficient as dextran in solution (static (1h,s) **Fig.2d**). In contrast, fibrin formed under flow exhibited reduced diffusion coefficients, which decreased further from 1 hour to 3 hours of flow. A statistical comparison of t_1/2_ and diffusion coefficients across different formation conditions in fibrin gels confirmed these trends (**Fig.2e-f**). The observed differences indicate slower diffusion of the 70 kDa dextran in fibrin formed under flow, with further reductions upon longer exposure to flow. This confirms that flow densifies the fibrin network and reduces pore size, thereby limiting the transport of certain enzymes to the network’s interior. Additionally, to assess the effect of incubation time on dextran transport, t_1/2_ and the diffusion coefficient were measured for static fibrin gels after 3 hours of incubation (**Fig.S8**). The formation time of static fibrin gels did not significantly affect dextran diffusion, as both the t_1/2_ and diffusion coefficient did not show significant differences in static fibrin gels formed for 1 hour, and in static fibrin gels formed for 3 hours. This highlights the dynamic effect of flow in densifying fibrin fibers over time and reducing the network pore size.

FRAP analysis provides a general understanding of how fast molecules diffuse within fibrin gels formed under different conditions. However, rapid diffusion does not necessarily ensure access to all fibrin molecules. Even at equilibrium, some fibrin molecules may be buried within the network or fiber interior, leading to reduced partitioning into fibrin-rich regions and decreased accessibility to lytic molecules. To assess the accessibility of fibrin-rich regions, pre-bleach images from the dextran FRAP experiments were segmented based on fibrin intensity, and the mean dextran fluorescence intensity was measured within the segmented regions of each image (**Fig.S9**). The smaller mean dextran intensity in fibrin fibers formed under flow indicates that dextran molecules partition less into fibrin-rich regions, even at equilibrium (**Fig.2g**). However, due to the diffraction limit of light, this analysis cannot distinguish between intra- or inter-fiber packing, particularly when fibers are spaced below the resolution limit. Therefore, we used FAM (5,6)-FITC as a tracer of intermolecular packing within fibrin fibers with fluorescence lifetime microscopy. As fibrin molecules pack more closely, the distance between FITC molecules may get closer, approaching the Förster distance (∼6 nm), where homo-FRET (Förster resonance energy transfer) and other dye-dye interactions become relevant. These interactions can lead to FITC self-quenching and a shorter fluorescence lifetime. A previous study using fibrinogen double-labeled with two fluorophores reported increased donor lifetime upon mechanical loading, which was attributed to fibrin unfolding and increased donor-acceptor separation in each molecule^71^.

To separate the intermolecular interactions from intramolecular effects, fibrinogen was labeled with approximately one FITC molecule per fibrinogen molecule, and only 5% labeled fibrinogen was used for fibrin clot formation. Since only 5% of fibrinogen molecules were labeled, the probability that two neighboring molecules within a protofibril both carried FITC was only 0.25%, strongly minimizing FITC-FITC interactions between adjacent molecules. Since fibrin molecules are ∼ 45 nm long (almost 8× the Förster distance), changes in FITC lifetime primarily reflect variations in the distance between FITC molecules on adjacent protofibrils which are spaced ∼13 nm apart^72^ (**Fig.2h**). This approach, however, neglects potential interactions between FITC molecules on adjacent fibrin fibers when the fibers are in the nanometer-scale proximity.

To validate the labeling strategy, we measured the FITC lifetime in mechanically stretched fibrin gels (**Fig.S10a**, supplementary Materials and Methods). Mechanical loading is known to increase protofibril alignment and packing within fibrin fibers^73^. As packing increases, FITC molecules come into closer proximity, increasing the likelihood of self-quenching. Consistent with this hypothesis, FITC lifetime decreased at 100% strain (**Fig.S10b**). FLIM of FITC-labeled fibrin formed under flow conditions (**Fig.2i**) showed an increase in fluorescence lifetime after 1 hour of flow which remained elevated after 3 hours of flow, compared to static fibrin clots formed under 1 hour of incubation (**Fig.2j**, distributions are shown in **Fig.S10c**). The longer lifetime indicates reduced intermolecular packing, suggesting increased spacing between neighboring protofibrils. Together, these findings suggest that although flow promotes the formation of a denser network of fibrin fibers, it may simultaneously reduce the density of protofibrils within the fiber structure.

### Flow alters the molecular structure of fibrin proteins in fibers

To investigate the effect of flow on the molecular structure of fibrin fibers, we performed BCARS spectral spectroscopy on fibrin gels across both the fingerprint region (Raman shift <1700 cm^-1^) and the CH region (Raman shift >2800 cm^-1^) (Fig.3a). Due to the high dimensionality of Raman spectra, PCA was used to reduce the dimensionality of data to the first two principal components, PC1 and PC2 (**Fig.S11a**). The distribution of data in PC2-PC1 space shifted from the second quadrant (static condition) to the fourth quadrant under flow (1 hour and 3 hours), indicating systematic changes in molecular features with flow and time. The loading scores identify the specific wavenumbers that contribute most strongly to these variations (**Fig.S11b**). In the fingerprint region, the Amide I band (∼1630−1700 cm^-1^) contributed strongly, while in the CH region, CH_2_ (∼2850 cm^-1^) and CH_3_ (∼2910−2065 cm^-1^) modes^44^ dominated. The Amide I band, corresponding to C=O stretching, is particularly sensitive to protein secondary structure^44^. To quantify changes in secondary structure motifs, the Amide I region was analyzed using Lorentzian fitting, where each structural component was represented by a peak with fixed FWHM and constrained center positions (**Table S1**, **Fig.3b**)^45,74^. The α-helix and β-sheet motifs were represented by two and one Lorentzian peaks, respectively. Comparison of the α-helix and β-sheet composition revealed a transition from the α-helix to the β-sheet under flow, which became more pronounced with longer exposure to flow (**Fig.3c**). This transition is consistent with previously observed fibrin response to external mechanical loading, experimentally and in computer simulations^37–39,46,75^. The content of random coil and turns did not significantly change under flow (**Fig.S11c**).

To interpret the results of experiments (Fig.3a − c), we turned to computational molecular modeling. We performed computational interrogation of the dynamic structural changes in fibrin monomer across a range of fibrin-based systems using MD simulations of atomic structural models of fibrin and Langevin simulations of simplified C_α_-atom models of fibrin structure (Materials and Methods). The fibrin monomer contains a central E nodule connected through the coiled-coil regions to the C-terminal β- and γ-nodules at both ends of the fibrin molecule (**Fig.3d**). Because the fibrin molecule contains two approximately homologous halves arranged about the central E region, we truncated the full fibrin molecule to generate a half-fibrin model (half-Fn). The reduced half-monomer representation has substantially lowered the computational cost and has enabled direct resolution of the force-induced conformational transitions in the fibrin molecule^51^. We carried out all-atom MD simulations in explicit solvent on the half-Fn system under hydrodynamic conditions mimicking the experimental flow conditions (**Fig.3e**). The half-Fn molecular fragment exhibited progressive loss of compact tertiary structure over time and underwent a series of unfolding transitions, starting in the coiled-coil regions (**Video S1**). Subsequently, the simulations revealed gradual extension and destabilization of the fibrin structure, consistent with a transition toward less ordered conformations. These results support the experimental findings by directly visualizing the solvent-flow-induced unfolding of individual fibrin monomers at the atomic level and demonstrate, in agreement with experiment, that fluid flow induces structural remodeling of fibrin molecules.

Because the MD simulations of the fibrin molecule in a solvent flow turned out to be computationally expensive, we next employed the all-atom MD simulations in conjunction with the SASA-based implicit-solvent framework^51,53,54^. By applying pulling forces to fibrin fragment half-Fn to mimic the action of the solvent flow described above, we explored whether the impact of hydrodynamic forces on fibrin molecular structure could be replicated by tensile pulling. To investigate the molecular basis underlying the dynamic transition from α-helices to β-sheets in fibrin, we performed pulling simulations on the half-Fn/des-αC fragment of fibrin (see Materials and Methods) by constraining the central region and pulling residues on the surface of β- and γ- nodules (**Fig.S2a and Table S2**). The pulling speeds ( 3 × 10^4^ − 10^6^µm/s) were used to approximate the experimental flow conditions (1500 s^-1^ shear rate) and geometry of the flow chamber (Fig.1b). The force-dependent conformational response reveals a non-linear structural transition. As shown in the 3D Ramachandran density maps (**Fig.S12a**), the α-helical populations progressively decrease with increasing force, with a pronounced reduction beyond ∼20 pN. In the same range (∼21–40 pN), the β-sheet populations increase sharply, indicating a force-induced α- to-β transition in the coiled-coil. At higher forces (>40 pN), both the α-helix and β-sheet populations decline, and the fibrin monomer becomes largely unstructured at ∼100-pN force. Secondary structure analysis (**Fig.S12b**) shows the α-helix relative content dropping from ∼0.4 to less than 0.1 in the 10–25 pN force range, while the β-sheet relative content increases from ∼0.2 to ∼0.4 in the same force range, followed by a gradual decrease to ∼0.2 at higher forces. These results illuminate a force-dependent pathway featuring transient β-rich intermediate fibrin structures populated prior to global unfolding transitions under high mechanical load.

### Flow-induced remodeling of fibrin reduces Plg and tPA binding

For Plg activation, both Plg and tPA bind to fibrin, forming a ternary complex^76^ with Plg binding primarily to residues Aα148−160, and tPA binding primarily to residues γ312−324. To assess the Plg binding, fibrin fibers were incubated with FITC-labeled Plg and subsequently imaged for fluorescence intensity. The ratio of Plg fluorescence intensity to fibrin intensity within fibrin-positive regions (Plg/Fn) was used as a metric for relative binding under different conditions (**Fig.4a**). The mean Plg/Fn ratio was significantly reduced in fibrin formed under flow compared with static conditions (**Fig.4b**). Plg binding sites appeared sensitive to flow, as even 1 hour of fibrin formation under flow drastically reduced the Plg/Fn ratio. However, extending the flow from 1 to 3 hours did not result in a further significant reduction, suggesting that most flow-induced alterations to the binding sites occur early during fibrin formation.

Using lower fibrinogen and thrombin concentrations and longer Plg incubation to see if the amount of Plg was limited, we found that the Plg/Fn ratio remained unchanged in static fibrin gels (**Fig.S13**). This suggests that the Plg concentration and incubation time were sufficient to achieve binding equilibrium. However, as the FRAP results show (Fig.2), fibrin formed under flow is denser and exhibits reduced molecular transport, which could limit the penetration of Plg and its binding within the network, even if Plg is available in excess. Therefore, it could artificially reduce the measured Plg/Fn ratio, independent of any actual disruption of fibrin-binding sites.

To address these confounding factors, a more conservative analysis was performed. Under static conditions, all fibrin-segmented pixels were included in the analysis. In contrast, for fibrin formed under flow, only the top 20% of segmented pixels with the highest Plg/Fn intensity ratios were analyzed, as these regions were expected to represent fibrin with the greatest accessibility to the surrounding solution and the strongest enzyme binding. The threshold was selected based on the mean intensity ratio of the top 0.5–10% of pixels to retain regions with high Plg binding while excluding dye puncta and artificially elevated ratios. This approach assumes that, despite the denser fibrin network under flow, a subset of fibrin molecules remains sufficiently exposed to achieve saturated binding, represented here by the top 20% of fibrin-rich pixels with the highest Plg/Fn ratios. By restricting the analysis to these regions under flow while including all fibrin pixels under static conditions, the analysis intentionally favored higher Plg/Fn ratios under flow, providing a stringent test of whether reduced binding persisted under conditions biased toward maximal binding. Despite this conservative comparison, the Plg/Fn ratio remained significantly lower under flow (**Fig.4c**), indicating that the reduced binding is unlikely to be explained by impaired transport or insufficient Plg availability and instead reflects disruption of Plg-binding sites.

To assess tPA binding to fibrin under flow, a similar strategy was employed. Because fibrin contains more binding sites for tPA than for Plg, a higher tPA concentration and longer incubation time were used to maximize binding-site occupancy. Although the Plg and tPA binding assays were performed under different enzyme concentrations, the results were independently validated through conservative binding analyses and/or dedicated control experiments. After tPA binding to fibrin, the fluorescent images were segmented based on the fibrin channel, and the tPA/Fn intensity ratio was used to quantify tPA binding (**Fig.4d**). Comparison of the tPA/Fn ratio across conditions showed no significant difference between static fibrin and fibrin formed under 1 hour of flow (**Fig.4e**), in contrast to the reduction observed for Plg binding. However, after 3 hours of flow, the tPA/Fn ratio decreased significantly, suggesting disruption of tPA-binding sites on fibrin. A more conservative analysis was also performed for this condition. Even when compared to the top 20% of fibrin pixels with the highest tPA/Fn ratio, under 3 hours of flow, static fibrin still exhibited significantly greater tPA binding (**Fig.4f**). These findings indicate that the reduced tPA binding after prolonged flow (3 hours) is attributable to disruption of tPA-binding sites rather than to potential confounding effects. Overall, the Plg and tPA binding assays demonstrate that high shear flow can disrupt Plg and tPA binding sites on fibrin, with more rapid and extensive disruption for Plg binding sites.

### MD simulations show force-induced disassembly of Aα and γ fibrinolytic binding sites in fibrin

We employed the all-atom MD simulations in conjunction with the SASA-based implicit-solvent framework^51,53,54^ to systematically explore the impact of hydrodynamic forces on binding sites of tPA and Plg. We mimicked the hydrodynamic conditions of shear stress by applying a time-dependent tensile perturbation to the fibrin structure. To investigate the molecular basis for the reduced Plg and tPA binding propensity observed under experimental hydrodynamic flow conditions, we focused on the half-Fn/des-αC fragment of fibrin (see Materials and Methods) using multiple pulling geometries and various directions of dynamic force application (displayed in **Figs.5a, b** and **Table S2**). This model contains the binding site for Plg (and tPA) in the Aα chain (residues K148−S160) and the cryptic binding site in the γ-nodule (residues F312−G324) for tPA (**Fig.5c**), which allows these binding sites’ specific molecular responses to mechanical loading to be probed and characterized directly. This approach enabled us to sample the conformational transitions in fibrin structure under a broader range of pulling vectors and force configurations relevant to the under-flow conditions (Materials and Methods). The pulling speeds were selected to approximate the experimental flow conditions based on the geometry of the flow chamber and applied shear stress (see **Table S2** for more detail).

Across all pulling geometries and directions of force application, dynamic loading induced progressive elongation of both binding regions, reflected by increases in the size of the binding sites. A representative MD simulation trajectory illustrating the progressive disassembly of the Plg binding region in the Aα chain and the tPA binding site in the γ-nodule during force application is shown in **Video S2**. To quantify the size of each binding site, we measured the distance between the first residue and the last residue along the sequence. For example, the size of the Plg (and tPA) binding site on Aα-chain is the distance between the first residue K148 and the last residue S160. This region underwent a sharp extension at relatively low forces (∼15–40 pN) (**Fig.5d**), whereas the γ-chain segment F312–G324 (binding site for tPA) required substantially higher forces (∼30– 200 pN) to exhibit comparable extension (**Fig.5e**). The lower critical force range for the binding site disassembly observed for the Aα-chain segment is due to the conformationally flexible coiled-coil region. In contrast, the γ-chain segment is in the structurally folded γ-nodule and is stabilized by tertiary structure interactions, resulting in overall greater mechanical resistance to disassembly.

In addition to monitoring the expansion of the individual binding sites, we quantified the distance between the AαK148–S160 and γF312–G324 binding sites (inter-site distance) by measuring the separation between their centers of mass. Structural studies^77^ have shown that these two sequences are positioned in close spatial proximity within the D region, with an initial separation of approximately 45 Å, an arrangement thought to facilitate efficient Plg activation by bringing Plg and tPA into the same local environment. This inter-site distance increased dramatically at forces as low as ∼17 pN (**Fig.5f**). Depending on the pulling geometry, the center-of-mass separation approximately doubled (from ∼45 Å to ∼90 Å) over a force range of ∼10–40 pN. Notably, within this same force range, the AαK148–S160 and, in some cases, the γF312–G324 regions also underwent substantial force-induced structural distortion, indicating that loss of binding-site integrity occurs concurrently with increasing inter-site separation. Together, these changes are expected to impair efficient assembly of the tPA-Plg-fibrin ternary complex well before the maximum separations are reached.

Next, we carried out electrostatic surface map analysis of the Aα-chain segment K148–S160 (primary binding site for Plg) and γ-chain segment F312–G324 (binding site for tPA), which revealed pronounced force-dependent changes in binding-site topology (**Fig.5g,h**). Upon mechanical loading, both binding regions adopted altered conformations, which indicates impaired binding affinity for fibrinolytic enzymes. These findings support the existence of a molecular mechanism, according to which the hydrodynamic shear-induced mechanical deformation of fibrin monomers directly remodels the recognition sites and inhibits binding of fibrinolytic enzymes.

To connect the experimental single-molecule force responses with network-scale mechanics of fibrin, we also analyzed forced unfolding for the double-stranded fibrin oligomers FO4/4^67^ stabilized by the A:a and B:b knob-hole bonds and γ-γ crosslink. This oligomer network contains multiple fibrinolytic binding sites, including 14 sites in the Aα chain and 16 sites in the γ-nodules of its constituent fibrin monomers (**Fig.S14b**). The force-extension analysis revealed distinct mechanical thresholds for disruption of fibrinolytic binding regions, with the Aα-chain binding sites undergoing complete structural disassembly at ∼500 pN (**Fig.S14b**) and the γ-chain binding sites undergoing disassembly at ∼250 pN (**Fig.S14c**). By normalizing these force levels across the number of contributing binding sites, we obtained the effective averaged force scale for mechanical disassembly of the binding sites. Specifically, the mechanical work associated with unfolding of each binding site was calculated from the corresponding force-extension trajectories (**Fig.S14d**), and the total work contributions from all binding sites were summed. Next, by dividing the total work by the total number of binding sites, we estimated an effective force scale of ∼100 pN for the γ-chain binding site disassembly and ∼250 pN for the Aα-chain binding site disassembly.

## Discussion

The fibrin network is the structural scaffold for blood clots and pathological thrombi. Fibrin formation stabilizes clots, whereas its degradation is essential for clot dissolution. Importantly, fibrin structure is dynamic and can vary with the conditions under which the clots and thrombi form. For example, arterial and venous thrombosis occur under distinct hemodynamic conditions. The higher shear rates and shear stresses in arteries contribute to differences in thrombus formation mechanisms, composition, structure, and lysability compared with venous thrombi^78,79,9,14,13^. Because fibrin is a major determinant of thrombus structure and lysis, we hypothesized that arterial blood flow influences fibrin properties and contributes to the distinct characteristics of arterial and venous thrombi. However, investigating the direct effects of flow on fibrin remains challenging because forming fibrin from purified fibrinogen under flow conditions is difficult to achieve experimentally. Additionally, thrombin, the main physiological activator of fibrinogen, rapidly forms a fibrin network throughout the solution when mixed directly with fibrinogen.

To overcome these challenges, in this study, instead of mixing thrombin with fibrinogen, we coated a confined region with thrombin to locally activate flowing fibrinogen and form fibrin under flow. Combined with a parallel-plate flow chamber, this approach is relatively inexpensive, easy to implement, and compatible with structural analyses such as Raman spectroscopy and confocal microscopy, compared with a previously used system^24^. Using this platform, we found that arterial-like flow (shear rate: ∼1500 s^-1^) alters both the network structure of fibrin, producing denser and more aligned fibers, consistent with previous observations^24,80^.

Furthermore, flow-induced fibrin remodeling may impair fibrinolysis by reducing Plg and tPA binding, thereby limiting tPA-mediated Plg activation. Plg and tPA share lysine-dependent binding site situated at residues Aα148−160 with similar binding affinities (K_d_ ∼1 μM)^81,82^ while tPA binds to an additional lysine-independent binding site at residues γ312−324^83^. Furthermore, the αC domain (Aα392 − 610) contains strong noncompetitive binding sites both for tPA and Plg (K_d_∼ 16− 33 nM)^84^. While all of these binding sites are largely concealed in fibrinogen, the conversion of fibrinogen to fibrin induces structural changes thereby exposing these binding sites^81^. The second second-order rate constant of Glutamine Plg activation increases from 0.01 µM^-1^s^-1^ to 0.63 µM^-1^s^-1^ (or from 0.0019 µM^-1^s^-1^ to 0.1825 µM^-1^s^-1^ based on other reports^85^) in the presence of fibrinogen and fibrin, respectively^76^.

To better understand the experimental results, we turned to computational modeling. We employed the all-atom MD simulations of atomic structural models and Langevin simulations of simplified model of fibrin fragments to explore the impact of fluid flow on dynamic structural alterations in fibrin molecules and on the binding of tPA and Plg to fibrin. In these simulations, we mimicked hydrodynamic shear stress by applying a time-dependent pulling force to the fibrin structure^51,54,73^. To sample a broader range of force configurations relevant to fibrin under flow, the simulations were performed with multiple pulling directions relative to the fibrin axis. The pulling rates were chosen to approximate the experimental flow conditions. At a flow rate of 1.1 mL/min, the mean flow velocity across the chamber cross-section (0.145×3.6 mm) was ∼3.5×10^4^ μm/s, falling within the range of pulling speeds used in the simulations (10^4^ – 10^6^ μm/s). By correlating the results of experiments and simulations, we were able to arrive at a structure-based interpretation of the experimental results.

Several factors can alter the efficiency of Plg activation by tPA and the consequent fibrinolysis. First, fibrin must be accessible to both tPA and Plg. Although thrombi contain 60–90% water by mass^86^, facilitating enzyme transport, diffusion through the clot is also governed by fibrin pore size^87^. We observed that flow-induced fibrin formation under flow with ∼1500 s^-1^ shear rate resulted in fibrin densification, reduced 70 kDa dextran partitioning into the fibrin network, slowed dextran diffusion, and hence limited dextran access to fibrin fibers. This is consistent with the previous studies showing that plasma clots formed under flow have smaller pore sizes^80^.

At the sub-fiber level, this transport restriction is reinforced by the flow-dependent organization of fibrin molecules in protofibrils. Protofibrils have a diameter of ∼5 nm^51^ and an average spacing of ∼13 nm^72^, which is comparable to the dimensions of Plg and tPA (∼9 nm and ∼14 nm for Glu-Plg and Lys-Plg^77,88^ and 10−13 nm for tPA^89^). FLIM imaging using FITC-labeled fibrinogen (FITC undergoes self-quenching^31,90^) showed increased FITC lifetime in fibrin clots formed under flow. Larger lifetime and reduced self-quenching suggest that fibrin molecules are, on average, more widely spaced under flow than under static conditions. The increased intermolecular spacing may facilitate the transport of lytic enzymes within the fiber interior and improve access to fibrin binding sites.

Second, the tPA and Plg binding sites on fibrin molecules must remain intact for efficient fibrinolysis. Any factors that disrupt these binding sites can reduce Plg activation. Experimentally, flow reduced both Plg and tPA binding to fibrin, consistent with previous observations of reduced tPA binding following fibrin stretching^15^. Plg binding decreased after 1 hour of flow, whereas tPA binding remained unchanged until 3 hours of flow. These results suggest a differential mechanical sensitivity of the two binding systems, with Plg binding sites being disrupted earlier and more readily than those involved in tPA binding. TPA engages fibrin through both Aα148−160 and γ312−324 positions, with the latter providing the dominant contribution to binding^91^. The finger domain of tPA mediates a lysine-independent interaction with residues γ312−324 on fibrin, while its kringle-2 (K2) domain contributes lysine-dependent contacts that may overlap with Plg binding sites such as Aα148−160. Accordingly, tPA lacking the finger domain shows substantially reduced fibrin binding^92^ , while the removal of the K2 domain causes only a modest change in binding affinity (K_d_ changes from 0.05 μM to 0.13 μM for the wild-type and K2-deficient tPA, respectively^93^). Similarly, lysine analogs, such as ε-aminocaproic acid, only weakly perturb tPA- fibrin interaction^81^. Together with the greater sensitivity of Plg binding under flow, these observations suggest that the binding site γ312−324 is more resistant to flow-induced unfolding than the binding site Aα148−160. Consistently, in simulations of single-molecule fibrin fragments under controlled mechanical loading, the Aα148−60 region shows earlier destabilization at lower forces (∼15–40 pN), whereas the γ312−324 region remains stable until higher forces (∼30–200 pN). This monomer-level hierarchy is further modified at the oligomeric scale. Mechanical testing of double-stranded fibrin oligomers revealed that effective force partitioning is redistributed by strand coupling and crosslinking, leading to an apparent inversion in disassembly thresholds: ∼100 pN for γ312−324-associated interactions and ∼250 pN for Aα148−160-associated interactions. This is because, in the simulations, mechanical tension is redistributed across coupled strands, where unfolding in one strand relaxes local tension and shifts load to the neighboring strand, producing alternating disassembly pathways^67^. In addition, in the simulations, we did not explicitly describe tPA and Plg enzymes bound to fibrin polypeptide chains, which might provide additional stabilization of the tertiary tPA-Plg-Fn complex, and so these force values should be viewed as force estimates characterizing the mechanical stability of the tPA-Fn and Plg-Fn interactions.

This multiscale fibrin monomer-to-oligomer description is particularly relevant for interpretation of the experimental results, since the fibrinogen preparation used in the flow assays is partially cross-linked, with a heterogeneous degree of γ-γ crosslinking^73^. Consequently, the experimental system is expected to exhibit a mixed mechanical response, retaining aspects of the monomeric hierarchy at early deformation regimes while progressively transitioning toward network-like force redistribution under sustained loading. In this framework, the early reduction in Plg binding is consistent with the lower-force susceptibility of Aα148−160-associated interactions, whereas the later decrease in tPA binding reflects the higher-force regime in which network-level load redistribution and crosslink-assisted stabilization become increasingly dominant.

Third, Plg and tPA must remain in proximity when bound to fibrin. The average distance between Aα148−160 and γ312−324 is ∼45 Å, which is approximately 2−3 times shorter than the reported lengths of Glu-Plg and Lys-Plg or tPA^77,88,89^. Flow-induced unfolding and increased separation between these binding sites reduce Plg activation even if the sites themselves remain intact. Raman spectroscopy of fibrin clots formed under flow revealed an α-helix to β-sheet transition, previously observed under large mechanical strains or slow pulling rates^38,39,54^. The present modeling efforts revealed the α-to-β transition under substantially higher pulling rates (10^4^−10^6^ µm/s), which more closely approximate the experimental flow conditions used in this study. Hence, the results obtained demonstrate that this response persists under physiologically relevant mechanical loading regimes. Such unfolding not only disrupts the dominant Plg and tPA binding sites but also increases the separation between them, potentially reducing their colocalization on fibrin and thereby limiting Plg activation.

Overall, these factors mainly focus on the role of fibrin as a substrate for Plg activation, which is a prerequisite for Pln’s role in tPA-driven fibrin digestion. Certainly, additional factors, including changes in Pln binding and cleavage sites, Factor XIII cross-linking, and fibrin-platelet interactions, may also contribute to reduced fibrinolysis and help explain the limited efficacy of thrombolytic therapy.

## Conclusions

Fibrin degradation by Pln is the basis of thrombolytic therapy. This process depends on the binding of tissue Plasminogen activator and Plasminogen enzymes to fibrin. When fibrin forms under flow conditions, such as those found in arterial thrombosis, it undergoes dynamic network and molecular structural remodeling that reduces the tissue Plasminogen activator and Plasminogen binding to fibrin. Computational modeling fully supports these observations and uncovers structural mechanisms of force-induced disassembly of the binding sites for Plg and tPA. These changes may impair fibrinolysis and contribute to thrombolysis resistance and the limited therapeutic time window – two longstanding challenges in stroke treatment.

## Acknowledgements

We acknowledge support from the National Science Foundation through grants 2105175, 2235856, 2127925, 2046148, 2404334, an American Heart Association Grant # 25IPA1456706, the National Institutes of Health through grant DA060543, the Welch Foundation through grants F-2008- 20220331, and the Office of Naval Research through grant N00014-23-1-2575.

## Notes

### Competing Interest Statement

The authors have declared no competing interest.

### Summary of Updates

Updated corresponding authors Updated typo corrections

