## Supplementary Data for "Flow alters fibrin molecular and network structure and decreases binding of fibrinolytic enzymes"

This file contains:

Supplementary Materials and Methods

Supplementary Figures S1-S14

Supplementary Tables S1-S3

Supplementary Videos S1 and S2

### Supplementary Materials and Methods:

**Adjusting the viscosity of Tricine buffer with Methylcellulose (MC):**

The viscosity of Tricine buffer at different MC concentrations was measured using particle-tracking microrheology[1]. Briefly, 3 µm microspheres (09850, Polysciences) were added to the buffer, which was then loaded into a ~200 µm chamber formed between two coverslips. Samples were maintained at 37 °C during imaging using the transmission channel of confocal microscopy (FV3000, Olympus). Image sequences were analyzed using the TrackMate plugin in ImageJ[2,3] to extract particle trajectories (**Fig.S1a**). Mean squared displacement (MSD) and lag time were computed and fitted linearly (**Fig.S1b**). For good linear fits with $R^2 > 0.96$, indicating normal diffusion, viscosity was calculated using the Stokes-Einstein equation. A MC concentration of 0.25% (w/v) yielded a viscosity of ~3.2 cP (**Fig.S1c**), consistent with that of blood. Water viscosity in room temperature and 37 °C were also measured as reference, matching their known values (**Fig.S1d**).

**Turbidity measurements:**

Turbidity measurements were performed using a BioTek Cytation 5 plate reader. Fibrinogen (3 mg/mL) in Tricine buffer or Tricine buffer + 0.25% MC was mixed with thrombin (1 U/mL) in 96-well plates immediately before measurement. Additionally, for the surface-initiated activation, wells were plasma-cleaned and coated with 10 U/mL thrombin for 15 minutes. The fibrinogen solution (3 mg/mL in Tricine + 0.25% MC) was then added to the wells. Turbidity was measured and quantified as described previously[4,5]. Briefly, absorbance at 405 nm was recorded every 15 seconds for 2 hours. The light pathlength was estimated using the same volume of water and absorbance measurements at 900 nm and 977 nm.

**Fluorescence lifetime imaging microscopy (FLIM) on mechanically stretched fibrin fibers:**

To evaluate the effect of mechanical loading on FITC lifetime, a control experiment was performed by casting a solution containing 1 mg/mL fibrinogen and 1 U/mL thrombin onto a plasma-cleaned PDMS sheet over a 10 mm × 5 mm area. After incubation for 1 hour at 37°C, PDMS sheets were stretched by 5 mm or 10 mm, corresponding to 50% and 100% strain, respectively. The stretched PDMS sheets were then mounted onto a coverslip using binder clips to maintain the applied strain. Finally, the samples were inverted onto a second coverslip separated by spacers for fluorescence lifetime imaging.

**Other image analysis:**

All other image analyses were performed in ImageJ and/or MATLAB. Fibrin fiber orientation was quantified using the OrientationJ plugin in ImageJ[6]. For each frame, the angle distribution was normalized to the range [0, 1] (technical replicate). The mean value across all frames in each experiment was then used as a biological replicate.

For sequential fluorescence imaging, frames were corrected for pixel misregistration using a custom macro code in ImageJ. The centroids of TetraSpeck Fluorescent Microspheres (T7284, Invitrogen™) in both channels were used as registration references.

**Flow characterization in the parallel-plate flow chamber:**

The parallel-plate flow chamber used in the experiments had a rectangular cross-section with dimensions $h \times w \approx 0.145 \times 3.6$ mm. The wall shear rate at the top and bottom surfaces is given by

$$\dot{\gamma} = \frac{6Q}{h^2 w},$$

where $\dot{\gamma}$ and $Q$ are the shear rate and the volumetric flow rate, respectively[7]. At the experimental flow rate of 1.1 mL/min, the wall shear rate was $\dot{\gamma} \approx 1453$ $s^{-1}$. Assuming Newtonian behavior, the corresponding wall shear stress is

$$\tau = \mu \frac{6Q}{h^2 w},$$

where $\tau$ is the wall shear stress and $\mu$ is the dynamic viscosity of the perfusate. Therefore, since the zero-shear dynamic viscosity of the Tricine buffer containing 0.25% (w/v) MC was 3.2 cP (3.2 mPa·s), the wall shear stress is $\tau \approx 4.6$ Pa.

### Supplementary Figures:

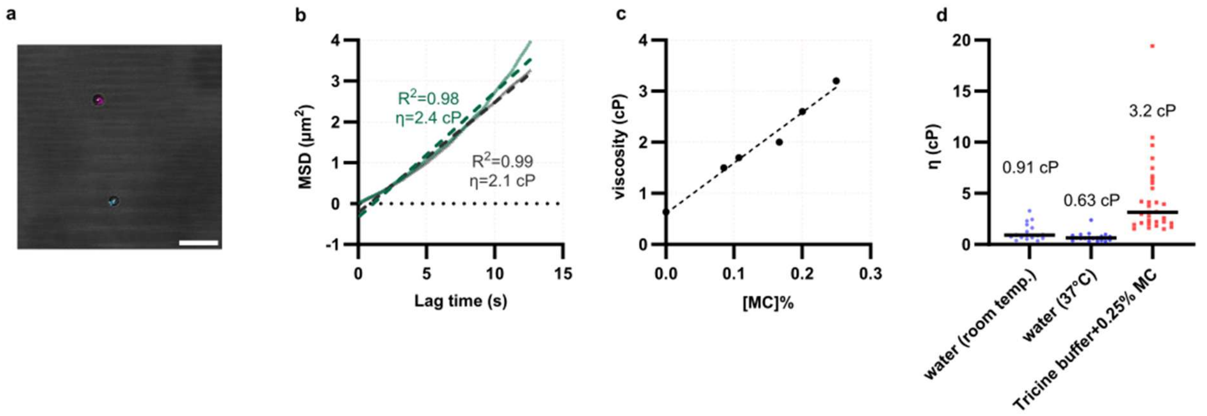


***Figure S1.*** *Viscosity measurements using particle-tracking microrheology.* ***a)*** *Brightfield image of two microspheres and their trajectories obtained using the TrackMate plugin in ImageJ (scale bar = 20 µm).* ***b)*** *Mean squared displacement (MSD) versus lag time for two representative microspheres, together with the corresponding linear fits and calculated viscosities.* ***c)*** *Measured viscosity as a function of methylcellulose (MC) concentration. The dashed line shows a fitted line.* ***d)*** *Measured viscosity of water at room temperature and 37 °C, and of Tricine buffer containing 0.25% (w/v) MC. Numbers indicate the corresponding median viscosities.*

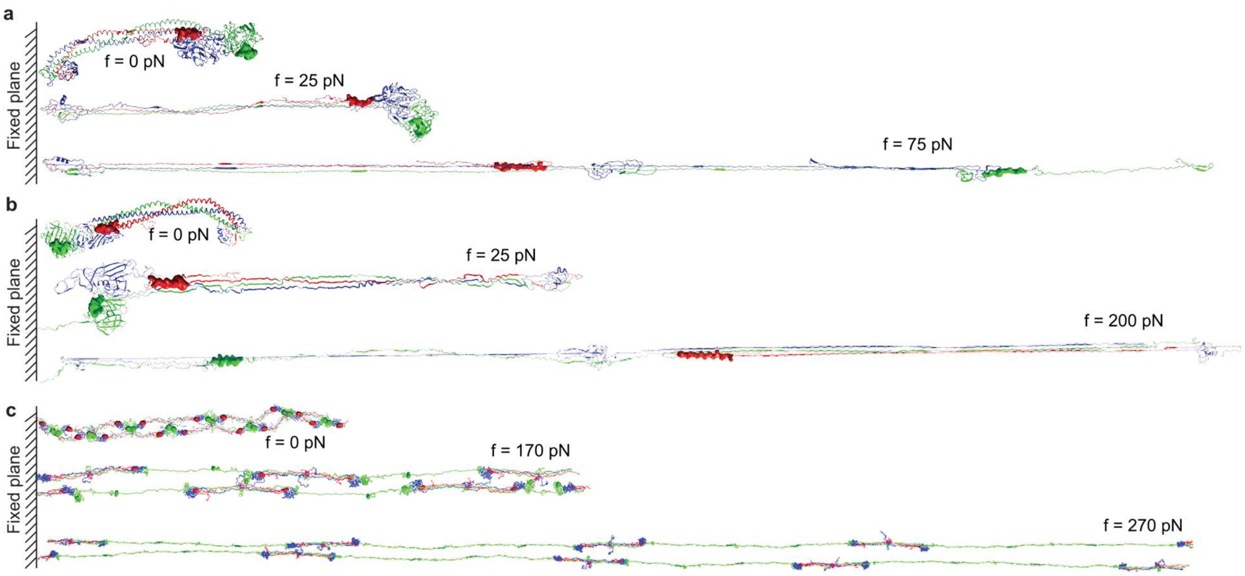


***Figure S2.*** *Force-induced unfolding transitions in fibrin structural fragments in silico.* ***a)*** *Representative snapshots from the dynamic force-ramp simulations for the half-Fb/des-αC system with the central E region fixed and pulling force applied to the β- and γ-nodules observed at 0, 25, and 75 pN.* ***b)*** *Representative snapshots from the dynamic force-ramp simulations for the Half-Fb/des-αC system with the constrained β- and γ-nodules and with the pulling applied to the central E region observed at 0, 25, and 200 pN.* ***c)*** *Representative snapshots from the dynamic force-ramp simulations for the γ-γ crosslinked, double-stranded fibrin oligomer FO4/4, with the pulling forces applied to the top and bottom rightmost fibrin monomers and with the top and bottom leftmost fibrin monomers constrained. In all panels, the binding sites for tPA and plasminogen are shown as molecular surfaces: Aα K148-S160 (for tPA and plasminogen) shown in red and γ F312-G324 (tPA) shown in green.*

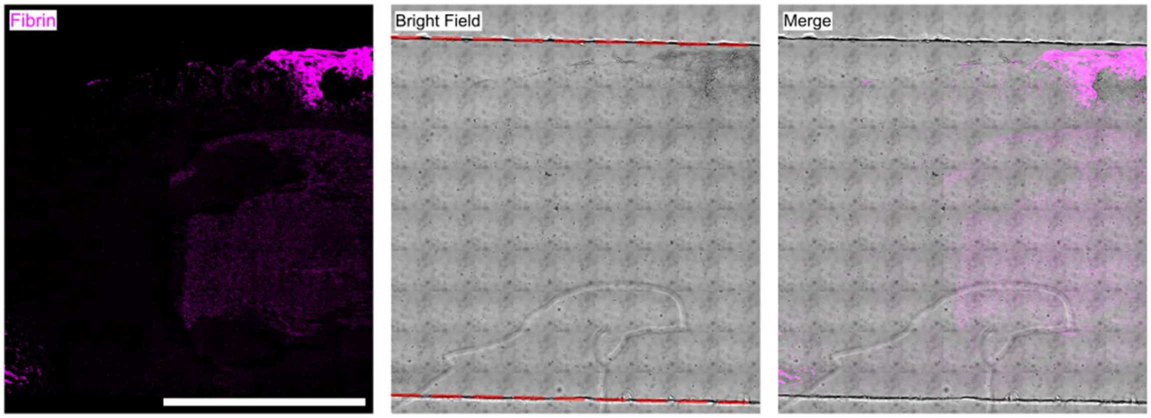


***Figure S3.*** *Fibrin formation inside the flow chamber under static conditions. Fibrin fluorescence intensity was contrast-adjusted for visualization purposes. Dashed red lines in the brightfield image (center) indicate the channel walls. Scale bar = 2 mm.*

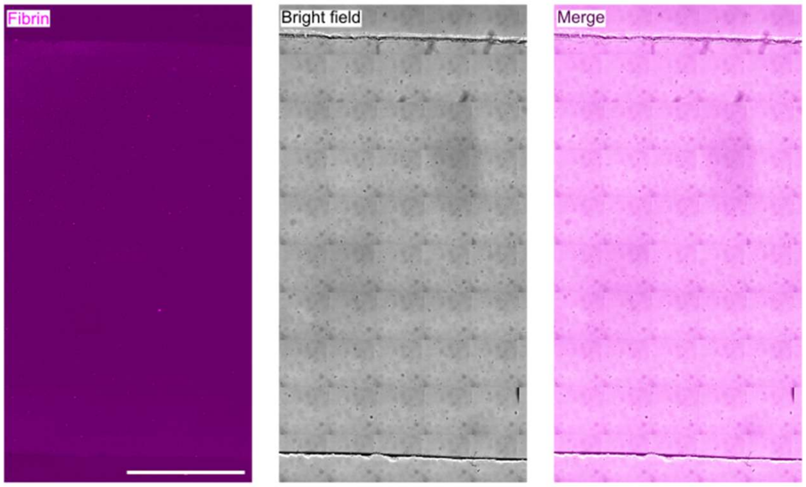


***Figure S4.*** *Representative image showing no fibrin formation after 3 hours of perfusion with fibrinogen solution in the absence of immobilized thrombin on the coverslip. Scale bar= 1mm.*

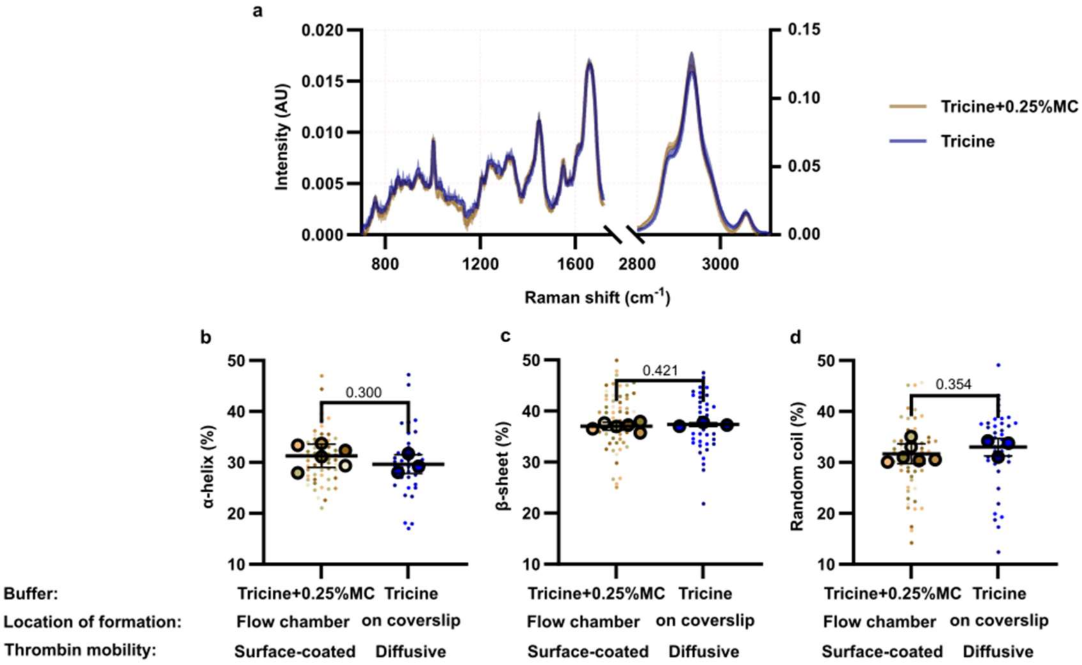


***Figure S5.*** *Comparison of fibrin gels formed under different buffer conditions (Tricine buffer vs. Tricine buffer + 0.25% MC), locations of formation (coverslip and flow chamber), and modes of thrombin delivery (surface-coated thrombin vs. soluble thrombin at 1 U/mL).* ***a)*** *Raman spectra of fibrin gels formed under the two conditions. The left y-axis corresponds to the fingerprint region (Raman shift < 1700 cm$^{-1}$), and the right y-axis corresponds to the CH region (Raman shift > 2800 cm$^{-1}$). Quantification of secondary structure content for* ***b)*** *α-helix,* ***c)*** *β-sheet, and* ***d)*** *random coil motifs. No significant differences were observed between the two groups.*

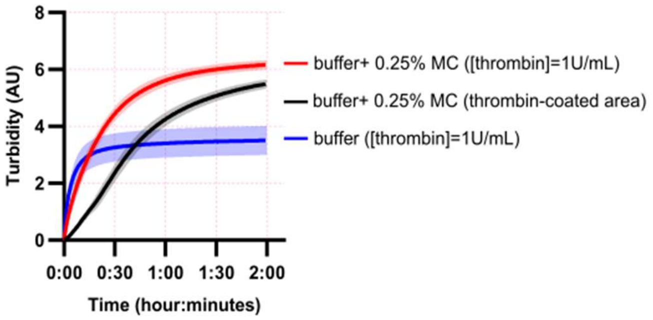


***Figure S6.*** *Turbidity measurement of fibrin formation under different conditions. Red and black curves were obtained in Tricine buffer containing 0.25% MC. Red and blue curves correspond to soluble thrombin, whereas black curve corresponds to surface-coated thrombin.*

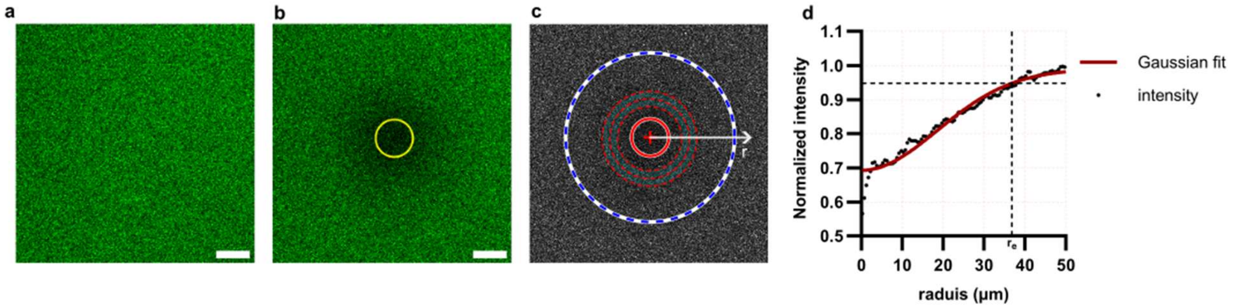


***Figure S7.*** *Intensity profile after photobleaching for calculation of the effective radius ($r_e$).* ***a)*** *Dextran fluorescence channel in a static (1h) fibrin gel before photobleaching.* ***b)*** *The same region after photobleaching. The yellow circle indicates the ROI.* ***c)*** *Normalized Dextran intensity, obtained by dividing the photobleached frame (b) by the pre-bleach image (a). The image is radially binned and averaged to obtain the radial intensity profile. The dashed blue circle indicates a radius of 50 μm.* ***d)*** *Radial intensity profile from the ROI center up to 50 μm, fitted with a Gaussian function to determine $r_e$. The scale bar is 20 μm.*

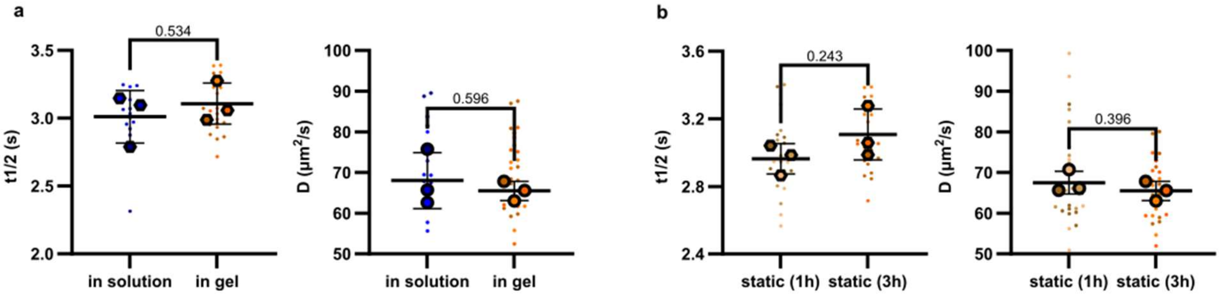


***Figure S8.*** *FRAP analysis of static fibrin gels formed after 3 hours of incubation.* ***a)*** *Comparison of $t_{1/2}$ and diffusion coefficient (D) between solution and fibrin gel.* ***b)*** *Comparison of $t_{1/2}$ and D in static fibrin gels after 1 hour and 3 hours of incubation.*

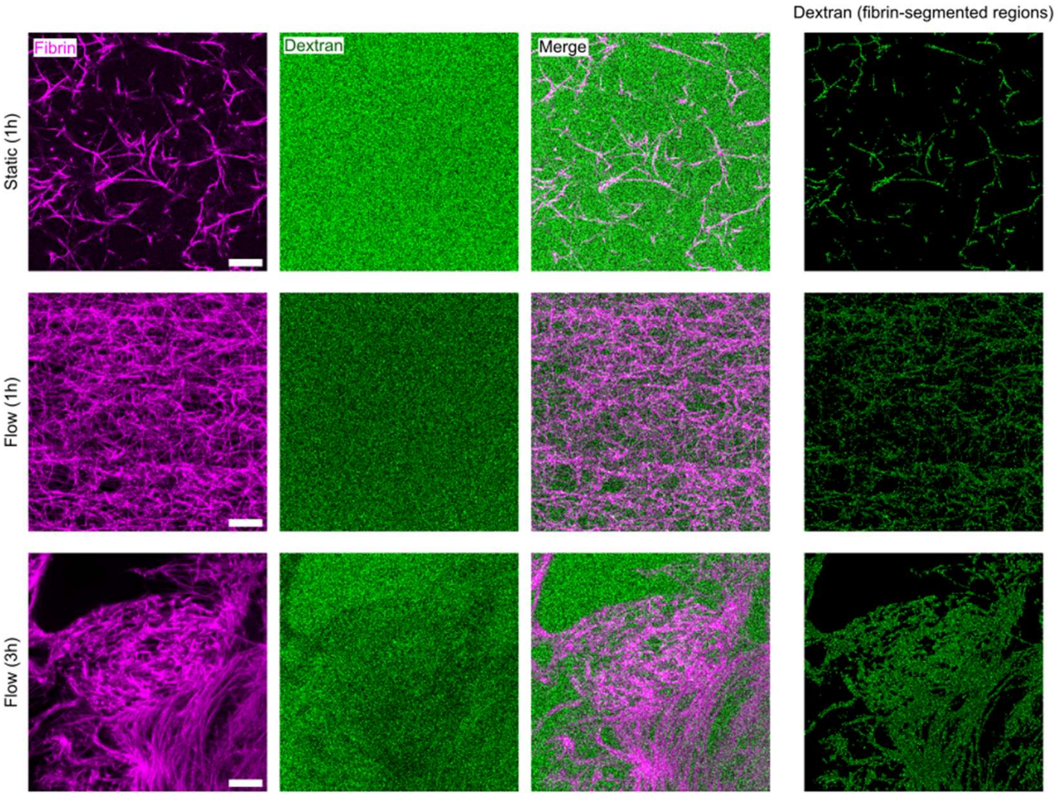


***Figure S9.*** *Dextran intensity in fibrin fibers across different groups. The pre-bleached image was segmented based on the fibrin channel, and the Dextran intensity was averaged within the segmented regions for each image. The Fibrin intensity range was adjusted for each image, but the Dextran channel intensity range was kept the same across all groups for visual comparison. The scale bar is 20 µm.*

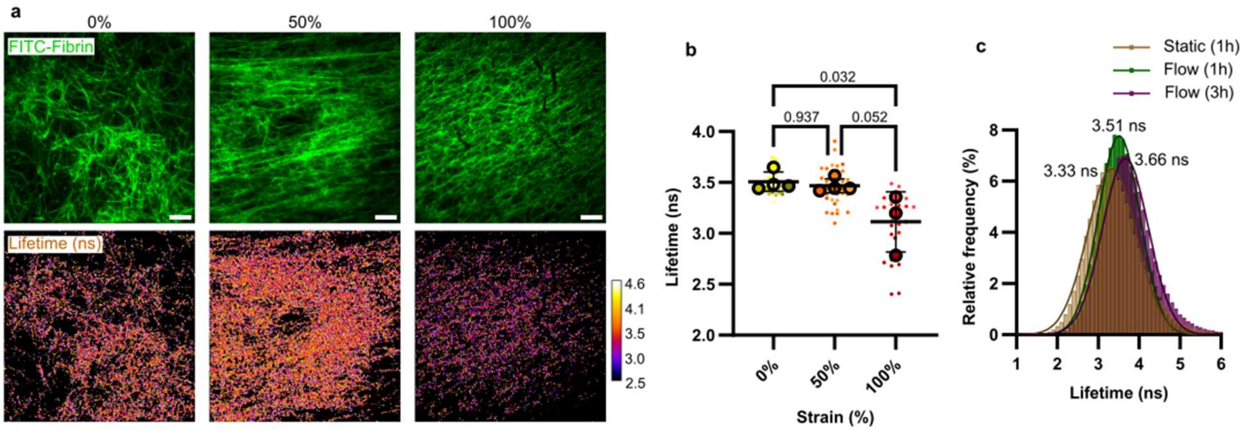


***Figure S10.*** *FITC lifetime changes under mechanical loading and under flow.* ***a)*** *Representative fluorescence intensity images of FITC-labeled fibrin gels under 0%, 50%, and 100% strain (top row), with the corresponding fluorescence lifetime maps (bottom row). Scale bar is 20 µm.* ***b)*** *Comparison of FITC lifetimes across strain conditions, demonstrating a decrease in lifetime under higher mechanical strain.* ***c)*** *Distribution of FITC lifetimes across all pixels under different flow conditions with fitted Gaussian curves. The values next to each peak indicate the lifetime at the maximum of the corresponding Gaussian fit. For clarity, only the lifetime range containing the peaks is shown, as lifetimes greater than 6 ns were rare and considered outliers.*

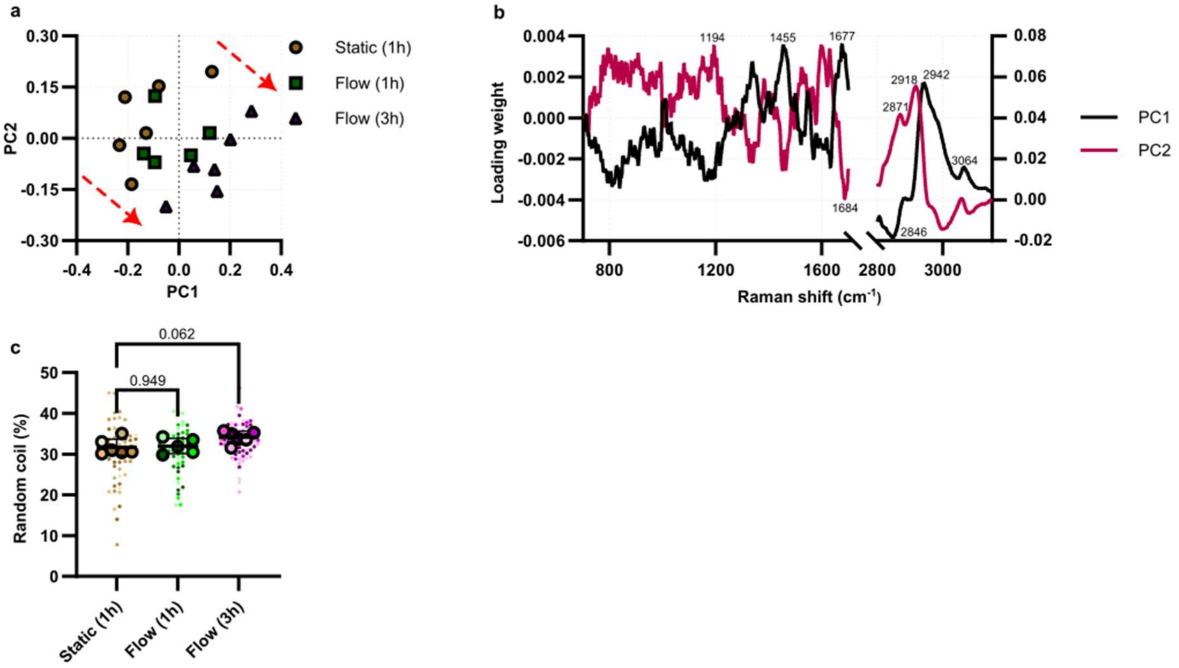


***Figure S11.*** *Raman analysis of fibrin gels formed under different conditions.* ***a)*** *PCA of fibrin spectra for different condition. Each point represents the PCA of the mean Raman spectrum for each experiment. The PCA scores shift from the second quadrant (static) to the fourth quadrant under 1 hour and 3 hours of flow.* ***b)*** *PCA loading scores highlighting the wavenumbers that contribute most to the separation. Selected peak positions are labeled. Loadings for the fingerprint region (Raman shift <1700 cm⁻¹) and the CH region (Raman shift >2800 cm⁻¹) are shown on the left and right axes, respectively.* ***c)*** *Comparison of random coil content across different fibrin formation conditions.*

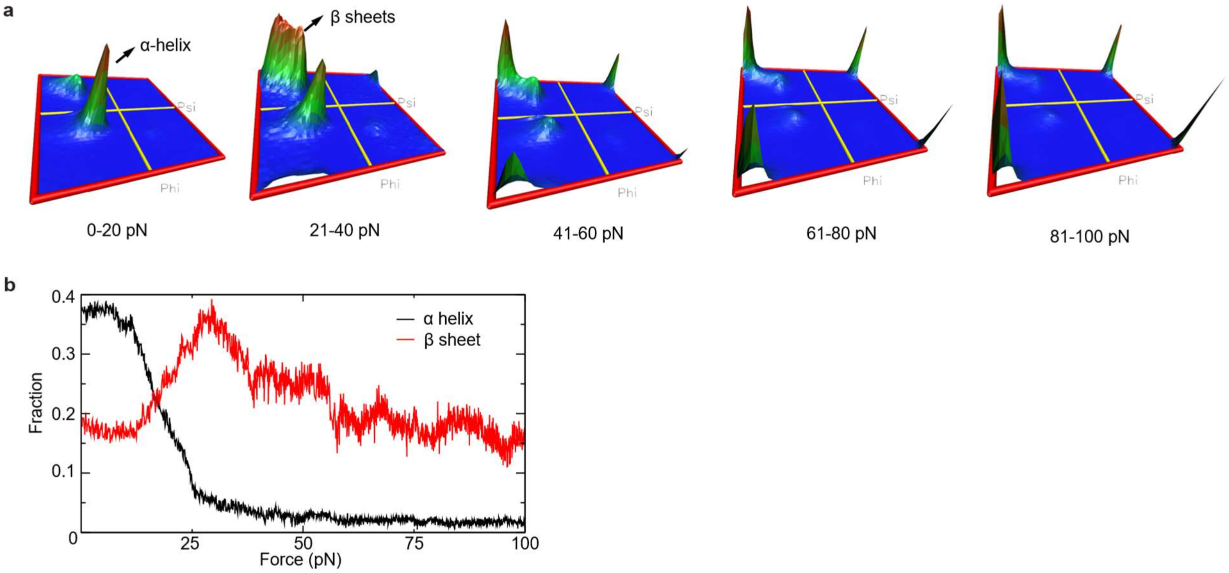


***Figure S12.*** *Force-induced secondary structure remodeling in half Fn/desαC during fibrin unfolding.* ***a)*** *Ramachandran density distribution at different force intervals during unfolding simulations (0–20 pN, 21–40 pN, 41–60 pN, 61–80 pN, and 81–100 pN).* ***b)*** *Fraction of α-helical and β-sheet secondary structure as a function of applied force during unfolding.*

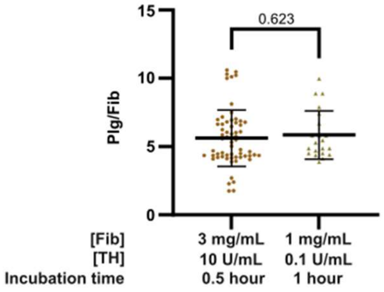


***Figure S13.*** *Comparison of two Plg-labeling scenarios with different fibrin content and similar Plg concentration in static conditions. In the second condition, fibrin was formed using lower fibrinogen concentration (1 mg/mL) on a thrombin-coated surface with 100× lower thrombin concentration (0.1 U/mL), followed by longer incubation with Plg (1 hour).*

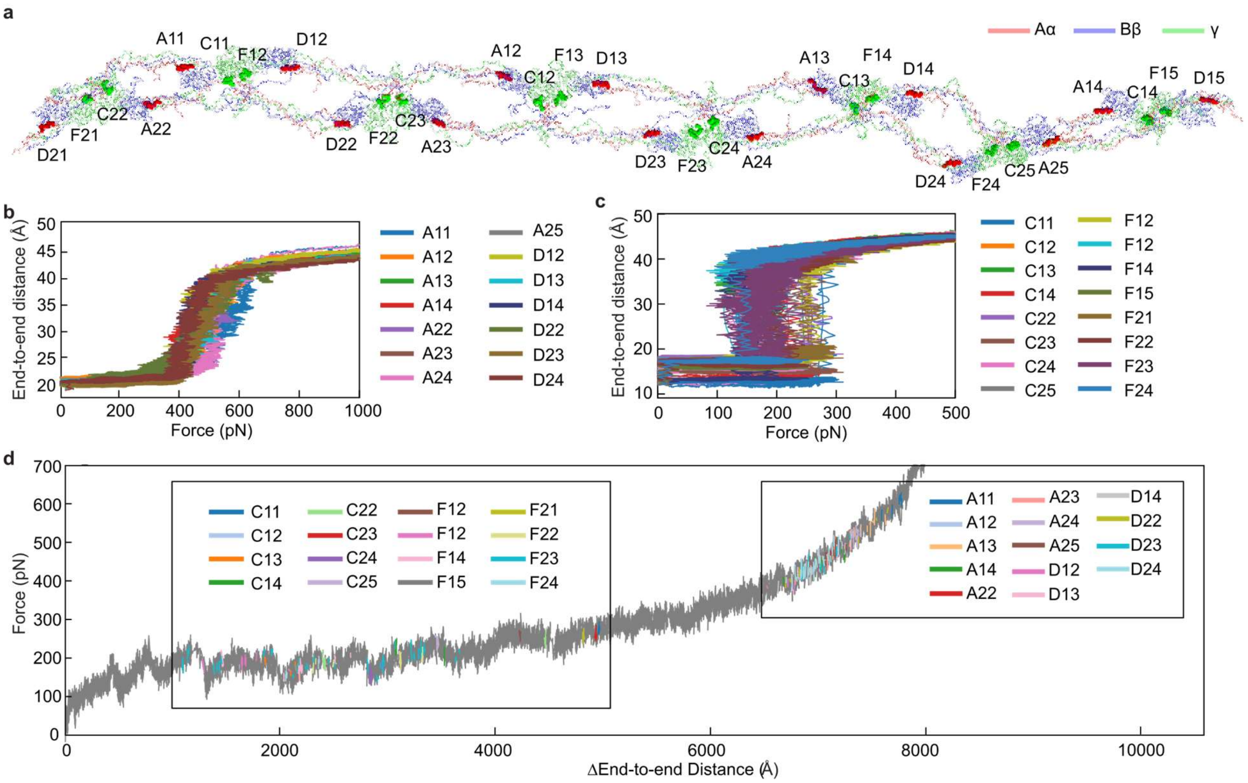


***Figure S14.*** *Mechanical disassembly of fibrinolytic binding sites in a double-stranded fibrin oligomer under tensile loading.* ***a)*** *Coarse-grained model of the γ-γ crosslinked double-stranded fibrin oligomer FO4/4 composed of fibrin monomers and containing multiple fibrinolytic binding sites, including 14 tPA- and plasminogen-binding sites in the Aα chains (red) and 16 tPA-binding sites in the γ-nodules (green).* ***b)*** *Distance between the $C_\alpha$-atoms spanning the Aα residues K148-S160 (size of the binding site for tPA and plasminogen) as a function of applied force.* ***c)*** *Distance between the $C_\alpha$-atoms spanning the γ chain residues F312-G324 (size of the binding site tPA) as a function of force.* ***d)*** *Global force-extension profile of the stretched fibrin oligomer, plotted as force versus the change in end-to-end distance. Colored segments highlight the regions of the trajectory corresponding to disassembly of individual fibrinolytic binding sites.*

### Supplementary Tables:

***Table S1.*** *Lorentzian fitting parameters for Amide I. Each motif is represented by a Lorentzian curve with a confined peak position and a fixed full width at half maximum (FWHM).*

| Secondary structure motif | Peak position range ($cm^{-1}$) | FWHM ($cm^{-1}$) |
|---|---|---|
| Tyrosine ring mode | 1595-1605 | 38 |
| Tyrosine ring mode | 1607-1617 | 32.7 |
| α-helix | 1638-1642 | 25.3 |
| α-helix | 1650-1654 | 17.5 |
| Random coil | 1658-1662 | 26.2 |
| β-sheet | 1671-1675 | 29 |
| β-turns | 1691-1695 | 24.4 |

***Table S2.*** *Force protocols used in Molecular Dynamics simulations of fibrin structure in explicit solvent. Listed are structural fragments used, constrained (or fixed) residues to maintain orientation during elongation and pulled residues (to which a pulling force was applied) in fibrin monomer, spring constant for constrained residues, as well as points of hydrodynamic force application in water molecules and direction of force application (relative to the fibrin longitudinal axis).*

| System | Structure portions | Constrained residues | Spring constant (kcal/mol/Å) | Bias target |
|---|---|---|---|---|
| Half-Fn | Central E domain | Aα17-47, Bβ59-79, γ1-22 | $10^4$ | Center-of-mass of water O-atoms |
| Half-Fn | β and γ Nodule | Aα358-610, Bβ364-457, γ325-410 | $10^4$ | Center-of-mass of water O-atoms |

***Table S3.*** *Force protocols used in Molecular Dynamics and Langevin simulations of fibrin structure. Listed are structural fragments of fibrin monomer, constrained (or fixed) residues, pulled residues (to which a pulling force was applied), and the values of pulling speed used in dynamic force ramp (see Methods in the main text).*

| **System** | **Structure portion** | **Constrained residues** | **Pulled residues** | **Pulling speed, μm/s** |
|---|---|---|---|---|
| Half-Fn/ des-αC | Central E domain: Set 1 | Bβ461, γ361 | Aα28, Aα36, Bβ65, γ8 | $10^6$ |
| Half-Fn /des-αC | Central E domain: Set 2 | Bβ461, γ361 | Aα28, Aα36, Bβ65, γ8 | $10^6$ |
| Half-Fn/ des-αC | β and γ Nodule: Set 1 | Aα28, Aα36, Bβ65, γ8 | Bβ386, 388, 390, 379; γ394, 158, 392, 188 | $10^5, 10^6$ |
| Half-Fn/ des-αC | β and γ Nodule: Set 2 | Aα28, Aα36, Bβ65, γ8 | Bβ353, 381, 386, 390, 379, 393, 372, 365; γ395, 159, 187, 162, 398, 400, 149, 407 | $3 \times 10^4$ |
| Half-Fn/ des-αC | β and γ Nodule: Set 3 | Aα28, Aα36, Bβ65, γ8 | Bβ334, 337, 361, 371; γ291, 295, 297, 373 | $10^5, 10^6$ |
| Half-Fn/ des-αC | β and γ Nodule: Set 4 | Aα28, Aα36, Bβ65, γ8 | Bβ361, 367, 371, 339, 352, 357, 343, 366; γ268, 266, 264, 235, 238, 279, 270, 291 | $3 \times 10^4$ |
| Half-Fn /des-αC | β and γ Nodule: Set 5 | Aα28, Aα36, Bβ65, γ8 | Bβ320, 318, 445, 448; γ329, 364, 360, 325 | $10^5, 10^6$ |
| Half-Fn/ des-αC | β and γ Nodule: Set 6 | Aα28, Aα36, Bβ65, γ8 | Bβ315, 319, 322, 348, 420, 442, 444; γ316, 322, 324, 329, 339, 341, 361, 364 | $3 \times 10^4$ |
| Fibrin oligomer[8] | Terminal pulling | Aα114, Bβ144, and γSer86 in the upper strand; Aα131, Bβ164, and γ105 in the lower strand of the leftmost fibrin monomer | Aα118, Bβ148, and γ90 in the upper strand; Aα127, Bβ160, and γ101 in the lower strand of the rightmost fibrin monomer | 1 |

### Supplementary Videos:

***Video S1.*** *Water-flow-induced unraveling of fibrin molecule. The movie shows an all-atom MD simulation for the half-Fn fibrin fragment performed in explicit solvent under hydrodynamic loading conditions. The αC domain and the β- and γ-nodules are held fixed while the solvent flow is directed toward the central E region. The trajectory illustrates the progressive loss of the tertiary contacts and secondary structure, starting in the coiled-coil region and followed by gradual extension of the fibrin molecule. Water molecules are shown explicitly, and the protein is displayed in the cartoon representation; the Aα chain colored red, the Bβ chain blue, and the γ chain green.*

***Video S2.*** *Force-induced disassembly of the fibrinolytic binding sites in fibrin molecule. The video shows an all-atom MD simulation of the forced unfolding for the half-Fn/des-αC fragment in implicit-solvent framework, illustrating conformational changes in fibrin structure under dynamic loading. The movie captures the progressive disassembly of the binding regions for plasminogen (Plg) and tissue plasminogen activator (tPA) in the Aα chain (residues K148–S160, highlighted in red surface representation) together with the tPA binding site in the γ-nodule (residues F312–G324, highlighted in green surface representation). A time-dependent force loading with a pulling speed of* $1 \times 10^5$ *μm/s was applied to the fibrin fragment to mimic hydrodynamic forces (see mathematical equation in the SI). Protein chains are shown in cartoon representation; the Aα chain colored red, the Bβ chain blue, and the γ chain green.*
